# A region-based method for causal mediation analysis of DNA methylation data

**DOI:** 10.1101/2020.11.03.366989

**Authors:** Qi Yan, Erick Forno, Juan C. Celedón, Wei Chen

**Affiliations:** Department of Obstetrics and Gynecology, Columbia University Irving Medical Center, New York, NY; Division of Pediatric Pulmonary Medicine, UPMC Children’s Hospital of Pittsburgh, University of Pittsburgh, Pittsburgh, PA; Department of Biostatistics, Graduate School of Public Health, University of Pittsburgh, Pittsburgh, PA

## Abstract

Exposure to environmental factors can affect DNA methylation at a CpG site or a genomic region, which can then affect an outcome. In other words, environmental effects on an outcome could be mediated by DNA methylation. To date, single CpG site-based mediation analysis has been employed extensively. More recently, however, there has been considerable interest on studying differentially methylated regions (DMRs), both because DMRs are more likely to have functional effects than single CpG sites and because testing DMRs reduces multiple testing. In this report, we propose a novel causal mediation approach under the counterfactual framework to test the significance of total, direct and indirect effects of predictors on response variable with a methylated region (MR) as the mediator (denoted as MR-Mediation). Functional linear transformation is used to reduce the possible high dimension of the CpG sites in a predefined methylated region and to account for their location information. In our simulation studies, MR-Mediation retained the desired Type I error rates for total, direct and indirect effect tests, for both continuous and binary outcomes. Furthermore, MR-Mediation had better power performance than testing mean methylation level as the mediator in most considered scenarios, especially for indirect effect (i.e., mediated effect) test, which could be more interesting than the other two effect tests. We further illustrate our proposed method by analyzing the methylation mediated effect of exposure to gun violence on total immunoglobulin E (IgE) or atopic asthma among participants in the Epigenetic Variation and Childhood Asthma in Puerto Ricans (EVA-PR) study.

## INTRODUCTION

Epigenetic studies are key to understanding regulatory mechanisms of gene expression. Of the available epigenetic markers, DNA methylation is the most stable and widely studied epigenetic modification.^1^ DNA methylation refers the addition of a methyl group at the 5′ location of cytosine nucleotides (C), with this modification occurring predominantly at Cs that are immediately followed by a guanine (G) in the 5′ to 3′ direction, denoted CpG. DNA methylation often affects gene transcription and has been associated with complex diseases^2–5^ and cancers.^6–10^

DNA methylation analysis has traditionally focused on individual CpG sites. Over the last few years, there has been great interest in developing region-based methods to detect differentially methylated regions (DMRs).^11, 12^ Such region-based approach is supported by strong correlation in DNA methylation levels across regions of the genome,^13, 14^ as well as by the fact that methylated regions such as CpG islands,^15^ CpG island shores,^16^ or generic 2-kb regions^17^ are more often linked to functionally relevant findings than single CpG sites. Statistically, if there are multiple causative CpG sites that have small individual effects, single-marker testing may have limited power to detect those weak signals. On the other hand, region-based approaches have higher power by combining the effects of multiple CpG sites. Moreover, region-based methods greatly reduce the burden of multiple testing in a genome-wide study.

DMR identification methods could be classified into two categories: 1. Supervised methods that first calculate the p-values for the association between each single CpG site and the outcome of interest, and then identify the regions with adjacent small p-values, and 2. Unsupervised methods that predefine the regions without using any outcome information, and then test the association between methylation level in the genomic regions and the outcome of interest.

Exposure to environmental factors can affect DNA methylation, which can then affect a disease or condition of interest. In other words, environmental effects on a disease or condition could be mediated by DNA methylation (Figure 1). Mediation analysis investigates the relationship between an exposure and an outcome, while examining how such exposure and outcome relate to the mediator. Although the Baron and Kenny’s “four steps” method^18^ has been the most commonly used approach to mediation analysis, such approach has been recently extended using a counterfactual framework^19–21^ to allow the presence of an interaction between the exposure and mediator, as well as to decompose the estimate of total effect (TE) into estimates of direct (DE) and indirect effects (IE). To date, single CpG site-based mediation analysis has been employed extensively. This approach has indeed shown that DNA methylation can mediate the effects of environmental exposure on complex traits,^22–24^ complex diseases^25–27^ and cancers.^28, 29^

**Figure 1.**
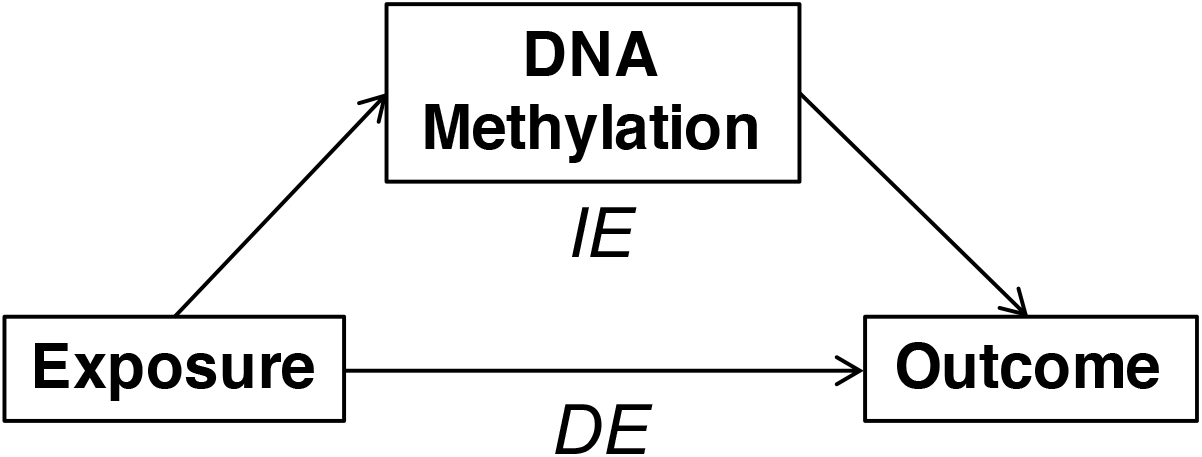
Mediation model with DNA methylation as the mediator.

In this study, we propose a novel causal mediation approach under the counterfactual framework to test the significance of total, direct, and indirect effects of DNA methylation in a genomic region on an outcome of interest. In this approach, a group of CpG sites from a predefined region are utilized as the mediator, and functional linear transformation^30^ is used to reduce the possible high dimensions in the region’s CpG sites and account for their location information. We denote the method as MR-Mediation. To evaluate the performance of the proposed MR-Mediation statistic, we first conduct extensive simulation studies to assess Type I error rates and power. We then illustrate our proposed method by analyzing whether DNA methylation mediates the association between exposure to gun violence and total immunoglobulin E (IgE) or atopic asthma among children participating in the Epigenetic Variation and Childhood Asthma in Puerto Ricans study (EVA-PR).^31^

## METHODS

### Functional linear transformation for region-based CpG sites

The idea is to linearly combine a few basis functions to fit a curve that can closely represent the methylation levels of the CpG sites in the region (e.g., an illustrative example is shown in Figure S1). Thus, the dimension can be reduced and the location information can be captured by the fitted curve. Specifically, we consider *n* subjects who have *m* CpG sites in a predefined region. The physical locations of the *m* CpG sites are normalized to a range of [0, 1] (i.e., 0 ≤ *l*_1_ … *l*_*m*_ ≤ 1). Thus, we use *M*_*i*_(*l*_*j*_) to denote the methylation level of a CpG site at *j*^*th*^ location from the *i*^*th*^ subject. We assume that the *n* × 1 vector of the continuous outcome **y** follows a linear regression model. For the *i*^*th*^ subject,

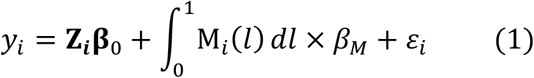

where **Z**_***i***_ is a 1 × *p* covariate vector, **β**_0_ is a *p* × 1 parameter vector (an intercept and *p* − 1 covariates), M_*i*_(*l*) is a function of CpG site locations and includes *m* CpG site values, *β*_*M*_ is the effect of M_*i*_(*l*), and *ε*_*i*_ is the random error. By using the ordinary linear smoother,^32^ we let

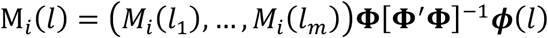

where (*M*_*i*_(*l*_1_), …, *M*_*i*_(*l*_*m*_)) is a 1 × *m* vector for *m* CpG sites in the region, ***ϕ***(*l*) is a *K*_*M*_ × 1 vector including *K*_*M*_ basis functions, and **Φ** is a *m* × *K*_*M*_ matrix containing the values of ***ϕ***(*l*) at *l*_1_ … *l*_*m*_. Thus, Formula (1) can be rewritten as

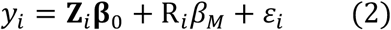

Where 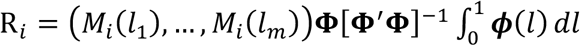 is a scalar. Therefore, R_*i*_ can be viewed as a summary of the methylation level in the region. In this study, we focus on cubic B-spline basis functions.

### Association between the mediator and the exposure variable

To test the proportions of the TE, DE and IE of the exposure on the outcome variables that are mediated by DNA methylation, we need to first make sure there exists an association between the mediator and exposure variables. To do so, we consider a linear regression model for *n* independent subjects,

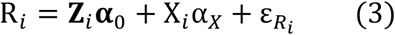

where R_*i*_ is a scalar representing the regional methylation level for the *i*^*th*^ subject, **Z**_***i***_ is a 1 × *p* covariate vector, **α**_0_ is a *p* × 1 parameter matrix (an intercept and *p* − 1 covariates), X*i* is a scalar for the exposure variable, α_*X*_ is the parameter for exposure, and 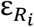 is a random error. In this study, we first test *H*_0_: α_*X*_ = 0, and then further test any methylation mediated effects between the exposure and outcome variables only when α_*X*_ ≠ 0.

### Counterfactual approach to the causal mediation model

The total effect (TE) from exposure to outcome can be decomposed into the indirect effect (IE) mediated through DNA methylation and the direct effect (DE) not mediated by DNA methylation, which could be mediated by other mechanisms or a direct link between exposure and outcome. For a continuous outcome, the mediation model for the *i*^*th*^ subject can be written as

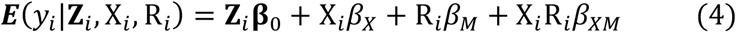

where **Z**_*i*_**β**_0_ is for the covariates, X_*i*_*β*_*X*_ is for the exposure variable, R_*i*_*β*_*M*_ is for the regional methylation level, and X_*i*_R_*i*_*β*_*XM*_ is for the interaction between the exposure variable and the regional methylation level. To use the counterfactual approach to the causal mediation model, four identifiability assumptions need to be satisfied: 1. no unmeasured confounding of the exposure and outcome relationship; 2. no unmeasured confounding of the mediator and outcome relationship; 3. no unmeasured confounding of the exposure and mediator relationship; 4. the exposure must not cause any known confounder of the mediator and outcome relationship.^33^ If all assumptions hold, the DE, IE and TE for change in exposure from level *x*_0_ to *x*_1_ are given by

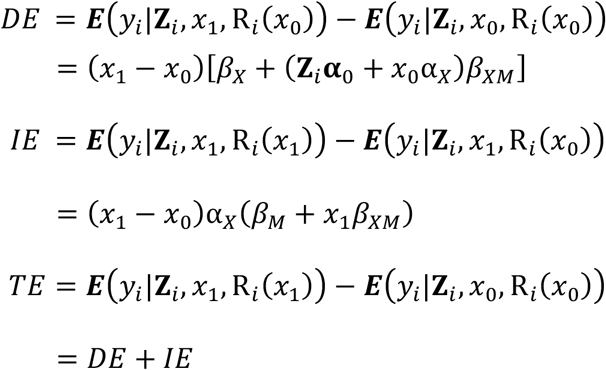

Since we only focus on the scenarios with α_*X*_ ≠ 0, conceivably, it can be shown that

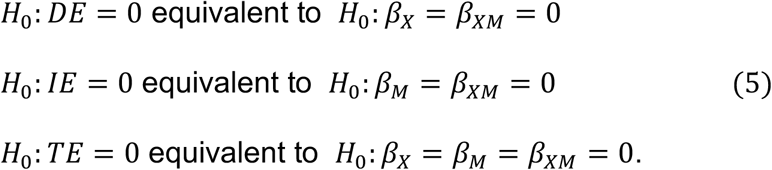

Thus, we can test *H*_0_: *DE* = 0 by a *F*-test with degrees of freedom (2, *n* − *p* − 3), *H*_0_: *IE* = 0 by a *F*-test with degrees of freedom (2, *n* − *p* − 3), and *H*_0_: *TE* = 0 by a *F*-test with degrees of freedom (3, *n* − *p* − 3).

Similarly, when the outcome is binary, the mediation model for the *i*^*th*^ subject can be written as

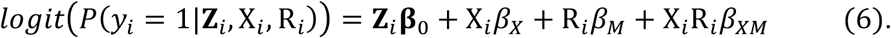

If all identifiability assumptions hold, the DE, IE and TE for change in exposure from level *x*_0_ to *x*_1_ are given by

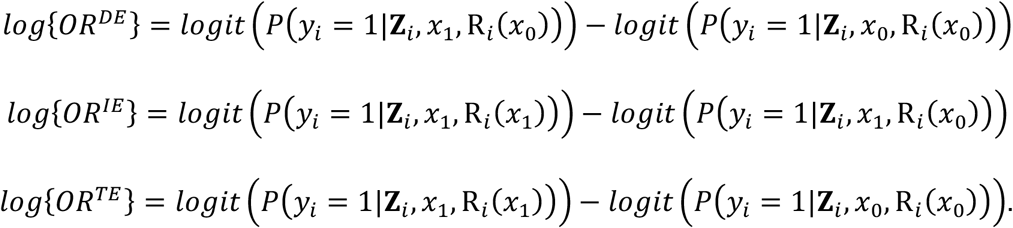

According to Valeri and VanderWeele (2013),^33^ a rare outcome assumption (i.e., low disease prevalence) is needed so as to have the odds ratio (OR) in the case-control design equivalent to the OR in the population, which is further approximately equivalent to the relative risk (RR) in the population. We also adopt the rare outcome assumption in this study. Then, the DE, IE and TE can be derived as

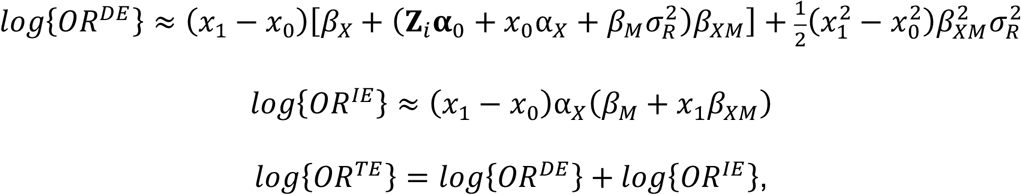

Where 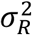 is the variance of random error from the mediator regression, Formula (3). The existence of 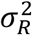 is due to 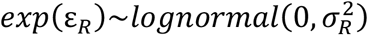. The null hypotheses for testing *DE* = 0, *IE* = 0 and *TE* = 0 for binary outcome are the same as given for continuous outcome. We can test *H*_0_: *DE* = 0 by a *χ*^2^-distributed Rao’s score statistic with 2 degrees of freedom, *H*_0_: *IE* = 0 with 2 degrees of freedom, and *H*_0_: *TE* = 0 with 3 degrees of freedom. Please note that the mediator regression, Formula (3), is run only for controls to account for the case-control design.^33^ In other words, because of the rare outcome assumption, the exposure distribution in the controls from the case-control design is approximately equivalent to the exposure distribution in the whole population.

### The Epigenetic Variation and Childhood Asthma in Puerto Ricans study (EVA-PR)

EVA-PR is a case-control study of asthma in subjects aged 9 to 20 years. Subject recruitment and study procedures for EVA-PR have been described elsewhere.^31^ Genome-wide DNA methylation was measured in 488 nasal epithelial samples using the Infinium HumanMethylation450 BeadChip arrays (Illumina, San Diego, CA). The preprocessing and quality control (QC) procedures were described in our prior study.^31^ After quality control, CpG sites with an overall mean β-value within [0.1, 0.9] were kept, leaving 227,901 CpG sites in the final dataset. Methylation M-value was calculated by log_2_(β-value/(1–β-value)). This methylation dataset was used in both the following simulation studies and in real data analysis.

The child’s lifetime exposure to gun violence was treated as the exposure variable in the following real data analysis. Lifetime exposure to gun violence was derived from the exposure to violence scale^34–36^ and analyzed as a binary variable (having heard gunshots at least twice vs. no more than once).^37^ Atopy was defined as an IgE ≥0.35 IU/mL to ≥1 of five common allergens (Der p 1, Bla g 2, Fel d 1, Can f 1, and Mus m 1). Asthma was defined as a physician’s diagnosis plus at least one episode of wheeze in the previous year. Atopic asthma was defined as the presence of both atopy and asthma, and thus treated as the binary outcome variable in the real data analysis; controls were non-asthmatic subjects. Total plasma IgE was treated as continuous outcome after log_10_-transformation.

### Simulation studies

We used the CpG sites from 488 nasal epithelial samples in EVA-PR for our simulation studies. The CpG sites were grouped into corresponding genes, and the kept CpG sites also needed to be within 5 kb distance from its nearest gene. The histogram plot (Figure S2) showed that most of the genes included less than 20 CpG sites, although some genes could have several hundred CpG sites. Thus, we selected different sizes of genes in our simulation studies. Specifically, we selected (1) 705 genes with 10 CpG sites; (2) 306 genes with 15 CpG sites; (3) 152 genes with 20 CpG sites; and (4) 21 genes with 40 CpG sites.

Furthermore, we need to choose the basis functions and the number of basis functions to reduce the dimension of each gene. To simulate the exposure **X** and outcome ***y*** variables, we used the same cubic (i.e., *order* = 4) B-spline basis functions for M_*i*_(*l*). We considered *K*_*M*_ = {4, …, 8} for all genes.

For each gene, we simulated one 488 × 1 vector for binary exposure **X** via the model:

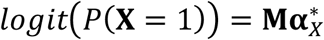

where **M** included 15% consecutive CpG sites covering the longest distance in the gene, each 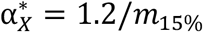 and *m*_1**5**%_ is the number of 15% CpG sites in the gene. The purpose is to make the exposure and the mediator (i.e., methylation) associated. Then, we can evaluate the power of testing α_*X*_ = 0 in Formula (3).

For each gene, we then simulated continuous and binary outcomes **y** for 488 samples via the models:

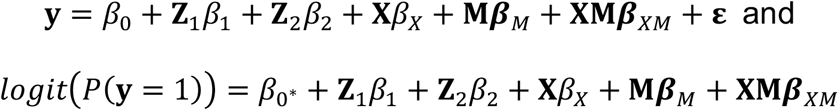

where **Z**_1_, **Z**_2_, **X** and **M** were the same as described above, **ε** was generated from standard normal distribution, *β*_0_ = 0.1, *β*_1_ = −0.1, *β*_2_ = 0.1 and *β*_0_∗ = −1. We considered (1) all *m*_15%_ effective CpG sites in the same direction (100%+) and (2) 50% effective CpG sites in the same direction (100%+) and (2) 50% effective CpG sites in one direction and the other 50% effective CpG sites in the opposite direction (50%+/50%-). We then considered eight settings for *β*_*X*_, ***β***_*E*_ and ***β***_*XE*_ to simulate ***y***, and these eight settings (Table 1) were used to evaluate Type I error rates and power of testing DE, IE and TE with the null hypotheses described in Formula (5). As shown in Table 1, Setting (1) was used to assess Type I error rates of testing DE, IE and TE; Setting (2) was used to assess Type I error rates of testing DE and power of testing IE and TE; Setting (3) was used to assess Type I error rates of testing IE and power of testing DE and TE; and Settings (4) ∼ (8) were used to assess power of testing DE, IE and TE. Please note that Setting (2) may not be the true null hypothesis for DE. Although *β*_*X*_ = 0, there could still be implicit **X** effect, because **X** and **M** are associated, and ***β***_*E*_ ≠ 0. Analogously, Setting (3) may not be the true null hypothesis for IE.

**Table 1.** The eight settings for *β*_*X*_, ***β***_*E*_ and ***β***_*XE*_ and their corresponding null hypothesis (*H*_0_) and alternative hypothesis (*H*_*a*_) for DE, IE and TE

|  | Settings<sup>*</sup> |  |  |  | DE | IE | TE |
| --- | --- | --- | --- | --- | --- | --- | --- |
|  | Continuous y |  | Binary y |  |  |  |  |
|  | 100%+ | 50%+/50%- | 100%+ | 50%+/50%- |  |  |  |
| (1) | $\beta_X = \beta_M = \beta_{XM} = 0$ | | $\beta_X = \beta_M = \beta_{XM} = 0$ | | $H_0$ | $H_0$ | $H_0$ |
| (2) | $\beta_X = \beta_{XM} = 0, \beta_M = 0.2$ | $\beta_X = \beta_{XM} = 0, \beta_M = 0.2/-0.2$ | $\beta_X = \beta_{XM} = 0, \beta_M = 0.4$ | $\beta_X = \beta_{XM} = 0, \beta_M = 0.4/-0.4$ | $H_0^{\#}$ | $H_a$ | $H_a$ |
| (3) | $\beta_M = \beta_{XM} = 0, \beta_X = 0.2$ | $\beta_M = \beta_{XM} = 0, \beta_X = 0.2/-0.2$ | $\beta_M = \beta_{XM} = 0, \beta_X = 0.4$ | $\beta_M = \beta_{XM} = 0, \beta_X = 0.4/-0.4$ | $H_a$ | $H_0^{\#}$ | $H_a$ |
| (4) | $\beta_X = \beta_M = 0, \beta_{XM} = 0.2$ | $\beta_X = \beta_M = 0, \beta_{XM} = 0.2/-0.2$ | $\beta_X = \beta_M = 0, \beta_{XM} = 0.4$ | $\beta_X = \beta_M = 0, \beta_{XM} = 0.4/-0.4$ | $H_a$ | $H_a$ | $H_a$ |
| (5) | $\beta_X = \beta_{XM} = 0.2, \beta_M = 0$ | $\beta_X = \beta_{XM} = 0.2/-0.2, \beta_M = 0$ | $\beta_X = \beta_{XM} = 0.4, \beta_M = 0$ | $\beta_X = \beta_{XM} = 0.4/-0.4, \beta_M = 0$ | $H_a$ | $H_a$ | $H_a$ |
| (6) | $\beta_M = \beta_{XM} = 0.2, \beta_X = 0$ | $\beta_M = \beta_{XM} = 0.2/-0.2, \beta_X = 0$ | $\beta_M = \beta_{XM} = 0.4, \beta_X = 0$ | $\beta_M = \beta_{XM} = 0.4/-0.4, \beta_X = 0$ | $H_a$ | $H_a$ | $H_a$ |
| (7) | $\beta_X = \beta_M = 0.2, \beta_{XM} = 0$ | $\beta_X = \beta_M = 0.2/-0.2, \beta_{XM} = 0$ | $\beta_X = \beta_M = 0.4, \beta_{XM} = 0$ | $\beta_X = \beta_M = 0.4/-0.4, \beta_{XM} = 0$ | $H_a$ | $H_a$ | $H_a$ |
| (8) | $\beta_X = \beta_M = \beta_{XM} = 0.2$ | $\beta_X = \beta_M = \beta_{XM} = 0.2/-0.2$ | $\beta_X = \beta_M = \beta_{XM} = 0.4$ | $\beta_X = \beta_M = \beta_{XM} = 0.4/-0.4$ | $H_a$ | $H_a$ | $H_a$ |

\* $\beta_M$ and $\beta_{XM}$ are vectors for 15% consecutive CpG sites. For 100%+, each element has the same assigned value. For 50%+/50%-, 50% elements have the same positive value and 50% elements have the same negative value. If the number of CpG sites is odd, one CpG site is assigned to have no effect.

For each gene, we simulated 10 sets of ***y*** for genes with 10 CpG sites (total is 705×10=7,050); 30 sets of **y** for genes with 15 CpG sites (total is 306×30=9,180); 50 sets of ***y*** for genes with 20 CpG sites (total is 152×50=7,600); and 400 sets of **y** for genes with 40 CpG sites (total is 21×400=8,400). Thus, we simulated 32,230 datasets for continuous outcome and another 32,230 datasets for binary outcome. After simulating the exposure **X** and outcome **y** variables, we used two approaches to conduct the analyses: (1) basis function fitted regional methylation level as the mediator (i.e., MR-Mediation) and (2) mean CpG methylation level as the mediator. We assessed the Type I error rates of testing DE, IE and TE only when the p-values of *α*_*X*_ were less than 0.05, and we evaluated the power at the significance level that the p-values of the corresponding effect (i.e., DE, IE or TE) and *α*_*X*_ were both less than 0.05.

## RESULTS

### Simulation of the Type I error rate

For both continuous and binary outcomes, both approaches retained the desired Type I error rates for DE, IE and TE across all considered gene sizes and basis numbers in Setting (1) (Figures 2, S3-S5 and S8-S10). Since Setting (2) might not be the true null hypothesis for DE, both MR-Mediation and the mean CpG methylation approach had inflated Type I error rates for scenarios with 100%+ effective CpG sites (Figure S6 and S11) and correct Type I error rates for scenarios with 50%+/50%-effective CpG sites (Figure S13 and S15). Although Setting (3) might not be the true null hypothesis for IE, both approaches retained the desired Type I error rates for all simulated conditions for both continuous and binary outcomes (Figures S7, S12, S14 and S16). Thus, both approaches are valid under their true null hypothesis.

**Figure 2.**
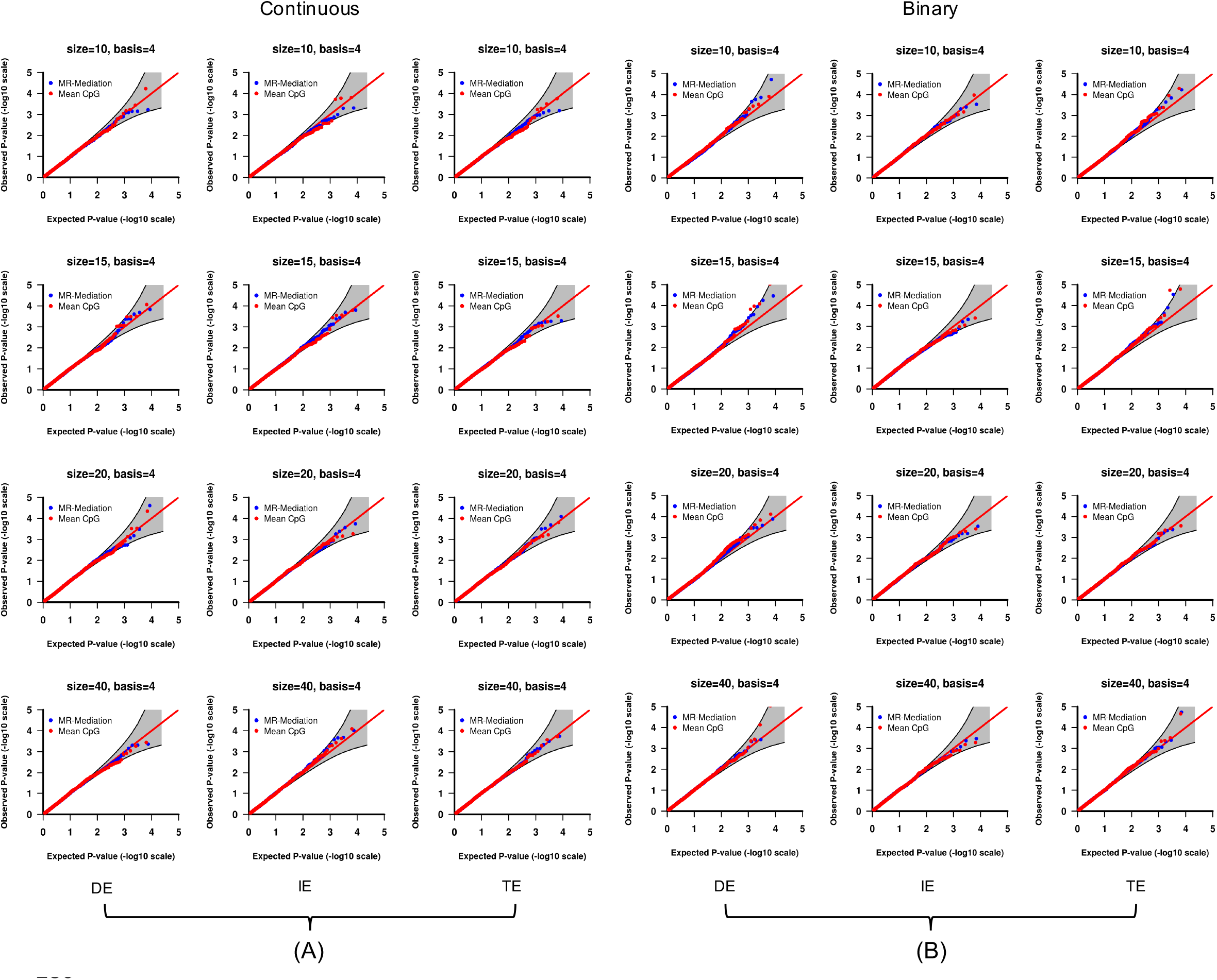
QQ plot of the *p*-values from Setting (1) with (A) continuous outcome and (B) binary outcome. The 1st column shows DE results, the 2nd column shows IE results and the 3rd column shows TE results. Each row shows one gene size (i.e., 10, 15, 20 or 40 CpG sites). Two approaches were compared: (1) MR-Mediation; and (2) Mean CpG methylation approach. A 95% pointwise confidence band (gray area) was computed under the assumption that the *p*-values were drawn independently from a uniform [0, 1] distribution.

### Simulation of power

In all considered scenarios, the power of MR-Mediation decreased as the basis number increased when the gene size was relatively small, and remained the similar power as the basis number increased when the gene size was large (Figures S17-S28). When the basis number was 4, MR-Mediation was the most powerful method in all scenarios for testing IE and more powerful than the mean CpG methylation method in almost all scenarios for testing TE and DE (Figures 3 and 4). In real data studies, one could be more interested in testing IE than TE and DE. Thus, MR-Mediation with the basis number equal to 4 is recommended.

**Figure 3.**
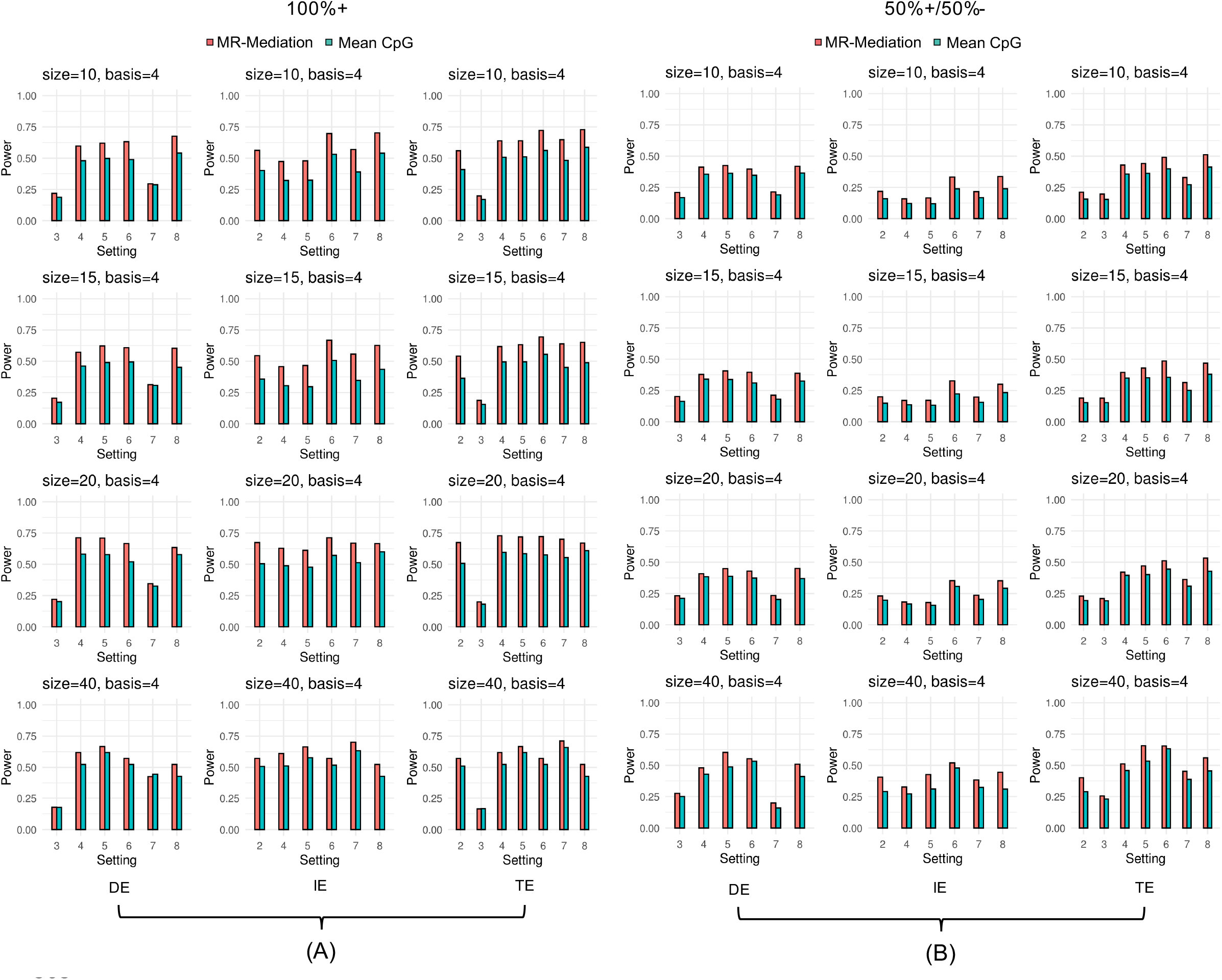
Power comparison for continuous outcome. (A) all positively effective CpG sites (100%+) and (B) 50% positively and 50% negatively effective CpG sites (50%+/50%-). The 1st column shows DE results, the 2nd column shows IE results and the 3rd column shows TE results. Each row shows one gene size (i.e., 10, 15, 20 or 40 CpG sites). Two approaches were compared: (1) MR-Mediation; and (2) Mean CpG methylation approach.

**Figure 4.**
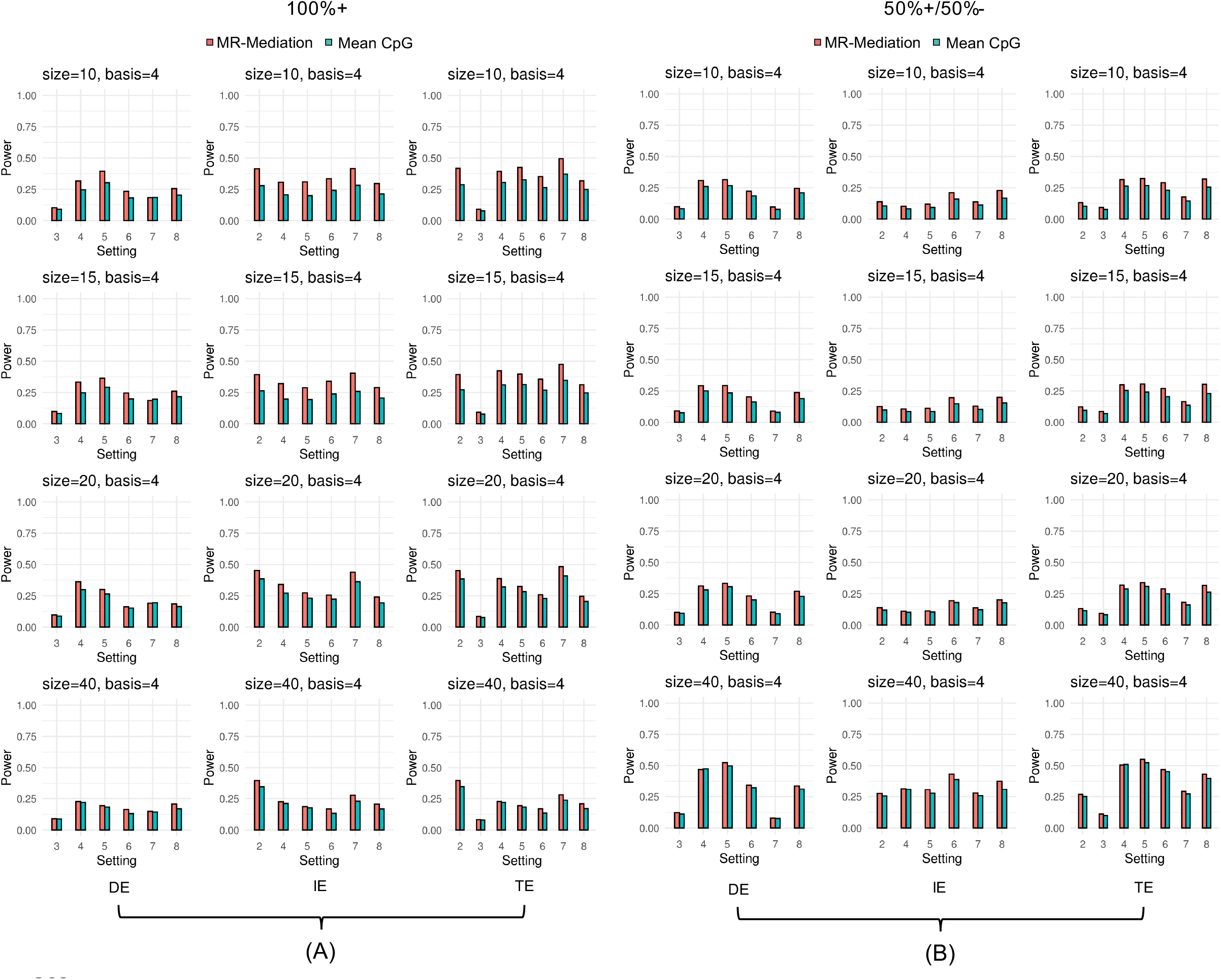
Power comparison for binary outcome. (A) all positively effective CpG sites (100%+) and (B) 50% positively and 50% negatively effective CpG sites (50%+/50%-). The 1st column shows DE results, the 2nd column shows IE results and the 3rd column shows TE results. Each row shows one gene size (i.e., 10, 15, 20 or 40 CpG sites). Two approaches were compared: (1) MR-Mediation; and (2) Mean CpG methylation approach. *Results in EVA-PR*

We used the proposed MR-Mediation method to study whether the association between exposure to gun violence and atopic asthma was mediated by DNA methylation, where atopic asthma is a binary outcome. In this study, we considered 5,720 genes including at least 10 CpG sties in 407 subjects without any missing covariates. Note that subjects who had asthma but no atopy were excluded from this analysis, as atopic asthma is the most common type of asthma in children. This analysis was adjusted for age, sex, annual household income (a measure of socioeconomic status), the top five principal components from genotypic data, methylation batch, and latent factors between exposure to gun violence and methylation, and between atopic asthma and methylation, estimated from *sva*.^38^ The mediator regression with four cubic B-spline basis functions, Formula (3), was run only for controls to account for the case-control design and we selected the top 20 genes for the further mediation analysis. We focused on the IE test to see if the association between exposure to gun violence and atopic asthma was mediated by methylation. We found that *CFD* on chromosome 19 was the top gene associated with exposure to gun violence (*P* = 1.93×10^−5^ and *FDR* = 0.1102, Table S1), although it did not reach the genome-wide significance level, and then it could mediate the effect of exposure to gun violence on atopic asthma risk (*P* = 3.09×10^−2^). The *CFD* gene has been reported to be associated with asthma.^39^ The expression of *CFD* can be reduced by IL-17A, which is required for development of airway hyperresponsiveness (a key intermediate phenotype of asthma). In addition, *TBC1D14* (*P* = 7.28×10^−3^) on chromosome 4, *FAM120B* (*P* = 3.33×10^−2^) on chromosome 6, *CRB1* (*P* = 1.75×10^−2^) on chromosome 10, *LOC339166* (*P* = 1.42×10^−2^) on chromosome 17, *ZGPAT* (*P* = 2.55×10^−3^) on chromosome 20, *MED16* (*P* = 2.05×10^−2^) on chromosome 19, *PLEKHG6* (*P* = 1.24×10^−3^) on chromosome 12, and *TNNI1* (*P* = 8.85×10^−3^) on chromosome 1 could also mediate the effect of exposure to gun violence on atopic asthma risk at the nominal level in the top 20 gene list.

We also applied MR-Mediation to study whether the association between exposure to gun violence and total IgE was mediated by DNA methylation, where total IgE is a continuous outcome. We also used 5,720 genes including at least 10 CpG sties in 473 subjects without any missing covariates. This analysis was adjusted for age, sex, annual household income, the top five principal components from genotypic data, methylation batch, and latent factors between exposure to gun violence and methylation, and between total IgE and methylation, estimated from *sva*.^38^ We selected the top 20 genes by *p*-values from the association between exposure to gun violence and gene-based methylation (Formula (3)) and conducted mediation analysis (Formula (4)). Again, we focused on the IE test to see if the association between exposure to gun violence and total IgE was mediated by methylation. The results showed that *PIP5K1C* (*P* = 4.33×10^−3^) on chromosome 19 could mediate the effect of exposure to gun violence on total IgE change at the nominal level in the top 20 gene list (Table S2).

## DISCUSSION

In this study, we developed a novel causal mediation approach, MR-Mediation, under the counterfactual framework to test the significance of DE, IE and TE. We implemented MR-Mediation in R (http://www.r-project.org) and the R package (https://cran.r-project.org/web/packages/MRmediation/index.html) is available. The major advantage of the counterfactual framework to mediation analysis is that it allows the decomposition of TE into DE and IE, even in models with nonlinearities and interactions. The counterfactual framework extends the widely used Baron and Kenny mediation approach by allowing an interaction term between exposure and mediator in the outcome regression.

We focused on testing the significance of DE, IE and TE instead of estimating their effects. For example, in IE, we test *H*_0_: *β*_*E*_ = *β*_*XE*_ = 0. When we reject the null hypothesis, it is possible that *β*_*E*_ = 0 and *β*_*XE*_ ≠ 0, which could not be known from the proposed test. Then the estimates of *β*_*E*_ may not be reasonable to be used in calculating IE. The parameters of covariates even need to be significant to calculate the estimates of DE and TE. In this approach, a group of CpG sites from a predefined region are utilized as the mediator, and the functional transformation is used to reduce the possible high dimension in the region-based CpG sites and account for their location information. The region-based analysis methods can improve power by combining weak signals and by reducing the multiple testing penalty. Users can use different ways to define regions of interest in addition to genes used in this study, such as, CpG islands and CpG shores.

For binary outcomes, we used the rare disease assumption, so that the OR in the case-control design is equal to the OR in the population, which is approximately equal to the RR in the population. When the disease is common in the population, the OR does not approximate the RR. In that case, a log link should be used in the generalized linear model for binary outcomes instead of a logit link, but the log link often fails to work with a binomial distribution. The Zou’s modified Poisson regression^40^ could be applied in this situation, although this is an approximate approach.

In the simulation studies, we show that MR-Mediation retains the correct Type I error rate for testing DE, IE and TE for both continuous and binary outcomes. In power comparison, we are more interested in IE, and MR-Mediation with basis=4 achieves better power than the mean CpG methylation approach. Our real data results show that MR-Mediation could help to identify potential methylation mediated effects. The genome-wide data analysis could be completed within an hour using one CPU.

Since MR-Mediation is a joint significance test, the exposure and regional methylation relationship and the regional methylation and outcome relationship are tested in two steps. It’s possible that CpG sites associated with the exposure and CpG sites associated with the outcome are different, which cannot be detected by the region-based mediator method. A further mediation analysis using single CpG sites may be needed. The single CpG site mediation function is also available in our R package. In addition, the summarized methylation level in the region can be viewed as the area under the curve. However, the areas could be still the same, even if curves are different. In future studies, we will consider to automatically partition the area into several sub-areas and each sub-area has a summarized methylation value. In such methods, ***α***_*X*_, ***β***_*E*_ and ***β***_*XE*_ are vectors instead of scalars as in this study. It’s possible that ***α***_*X*_(***β***_*E*_ + *x*_1_***β***_*XE*_) = 0, even if ***α***_*X*_ ≠ 0 and ***β***_*E*_ ≠ 0 or ***β***_*XE*_ ≠ 0. We will need to address this potential problem.

MR-Mediation can be extended to high dimensional predictors, such as multiple SNPs from a region, so as to investigate DNA methylation-mediated genetic effects on an outcome. These SNPs can be from the same region as the CpG sites. Such region-based approach on both SNPs and CpG sites can boost statistical power, because the region-based approach combines small individual effects and reduce multiple testing. Moreover, the region-based CpG sites can also be treated as predictors and gene expression can be treated as a mediator in order to understand the regulatory mechanisms of DNA methylation on gene expression that further affect diseases or traits.

## Supplemental Material

Supplemental data for this article can be accessed on the publisher’s website.

