## Supplemental Tables and Figures for "A region-based method for causal mediation analysis of DNA methylation data"

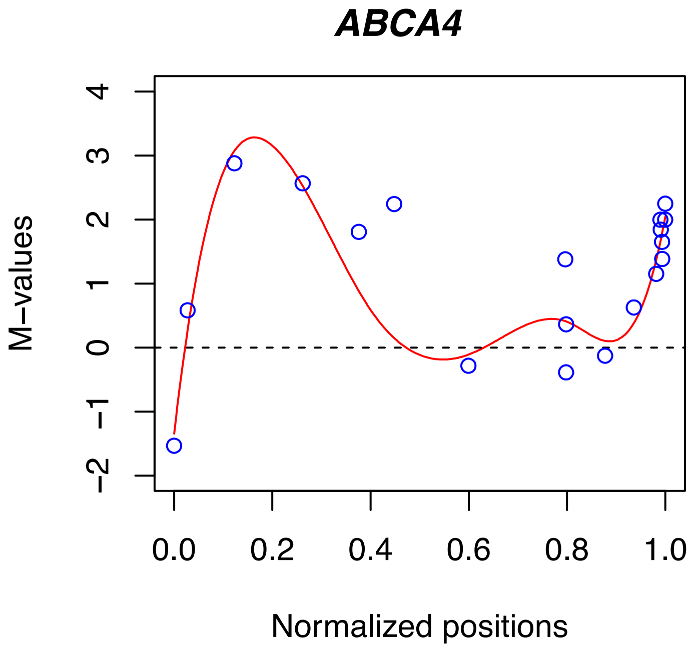


Figure S1. An illustrative example of CpG sites in gene *ABCA4* from one sample. The red curve is fitted by 8 cubic (order=4) B-spline basis functions.

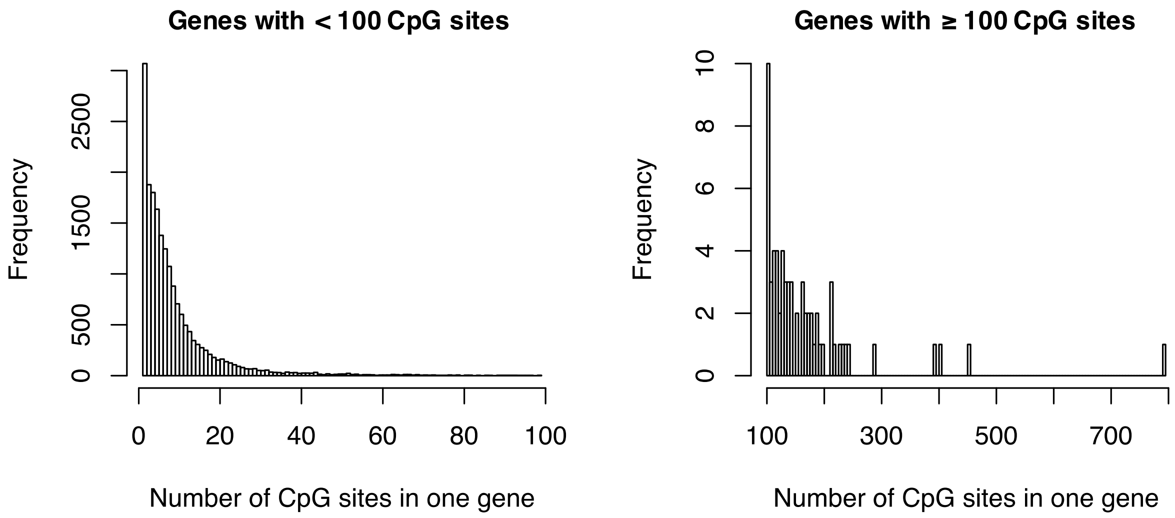


Figure S2. Histogram plots displaying the frequency of genes with different number of CpG sites included.

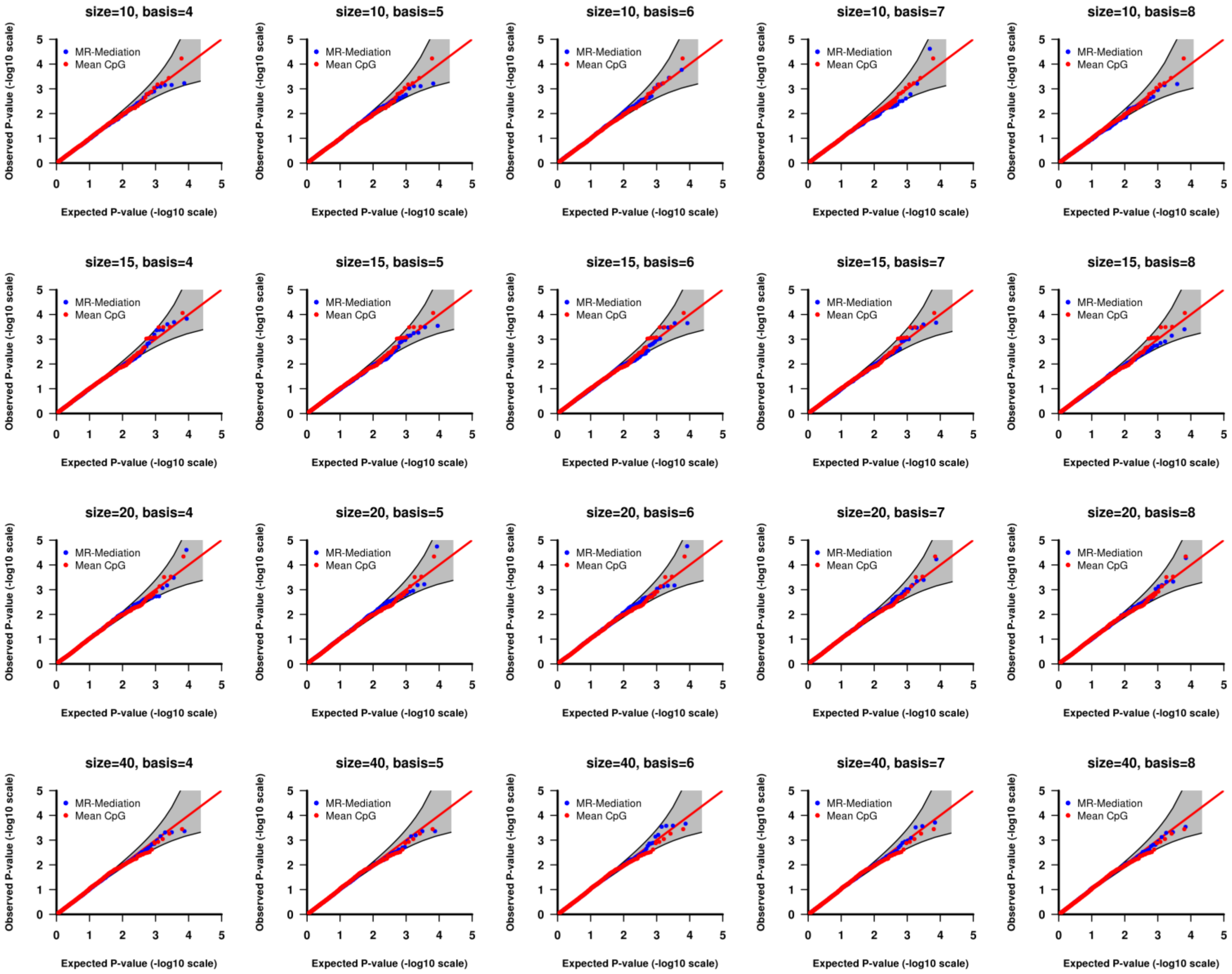


Figure S3. QQ plot of the *p*-values from Setting (1) DE tests with continuous outcome. Each row shows one gene size with different basis numbers. Two approaches were compared: (1) MR-Mediation; and (2) Mean CpG methylation approach. A 95% pointwise confidence band (gray area) was computed under the assumption that the *p*-values were drawn independently from a uniform [0, 1] distribution.

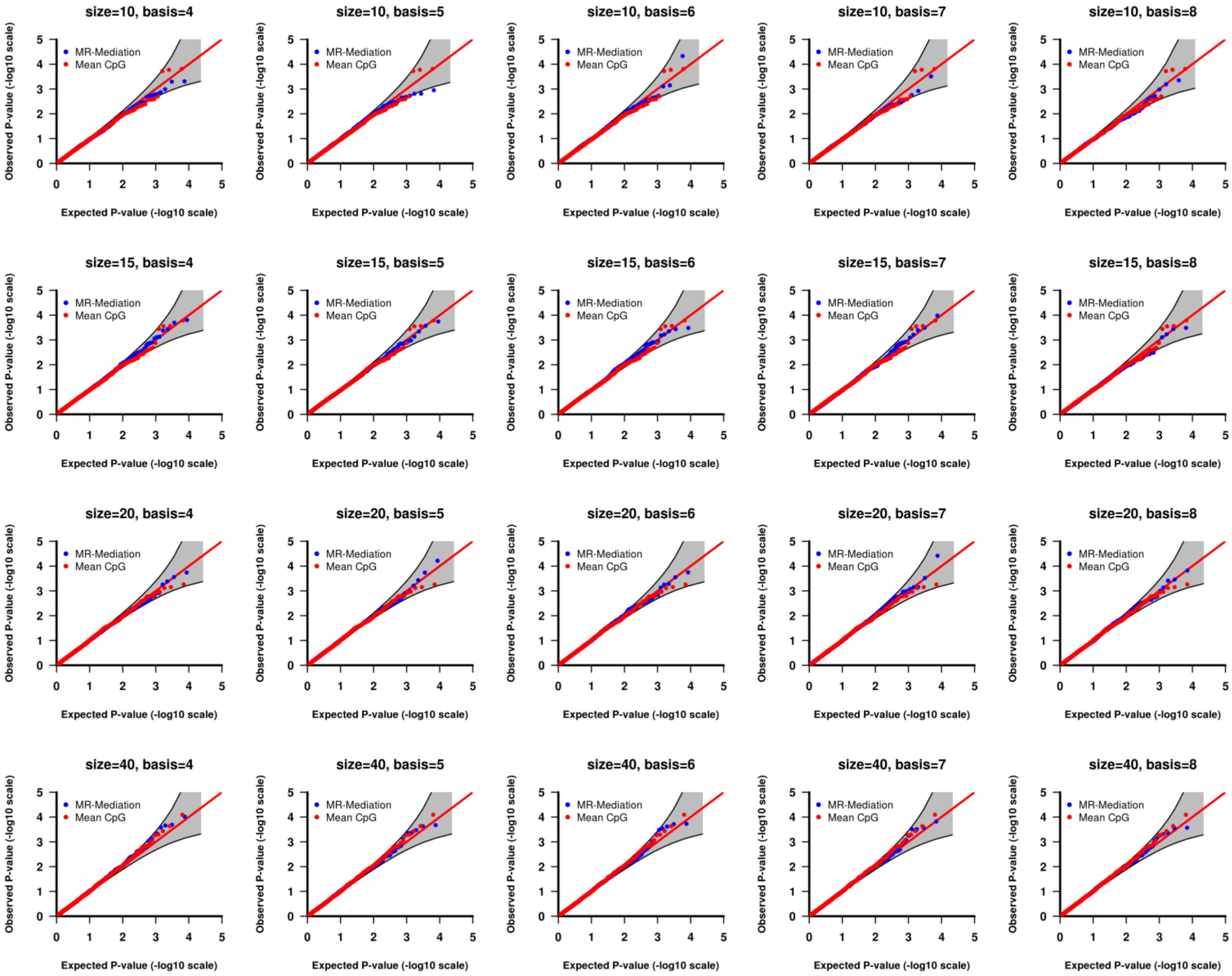


Figure S4. QQ plot of the *p*-values from Setting (1) IE tests with continuous outcome. Each row shows one gene size with different basis numbers. Two approaches were compared: (1) MR-Mediation; and (2) Mean CpG methylation approach. A 95% pointwise confidence band (gray area) was computed under the assumption that the *p*-values were drawn independently from a uniform [0, 1] distribution.

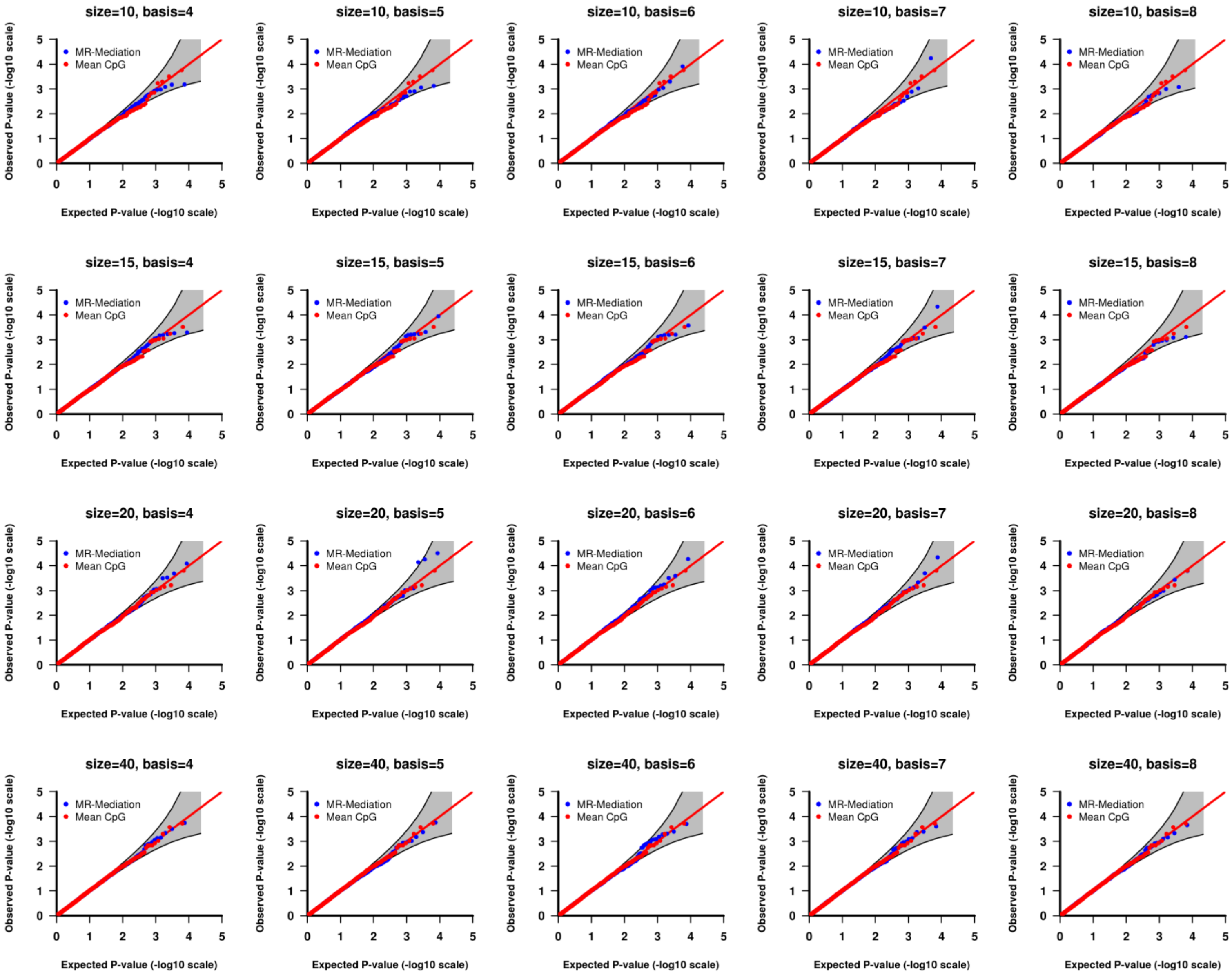


Figure S5. QQ plot of the *p*-values from Setting (1) TE tests with continuous outcome. Each row shows one gene size with different basis numbers. Two approaches were compared: (1) MR-Mediation; and (2) Mean CpG methylation approach. A 95% pointwise confidence band (gray area) was computed under the assumption that the *p*-values were drawn independently from a uniform [0, 1] distribution.

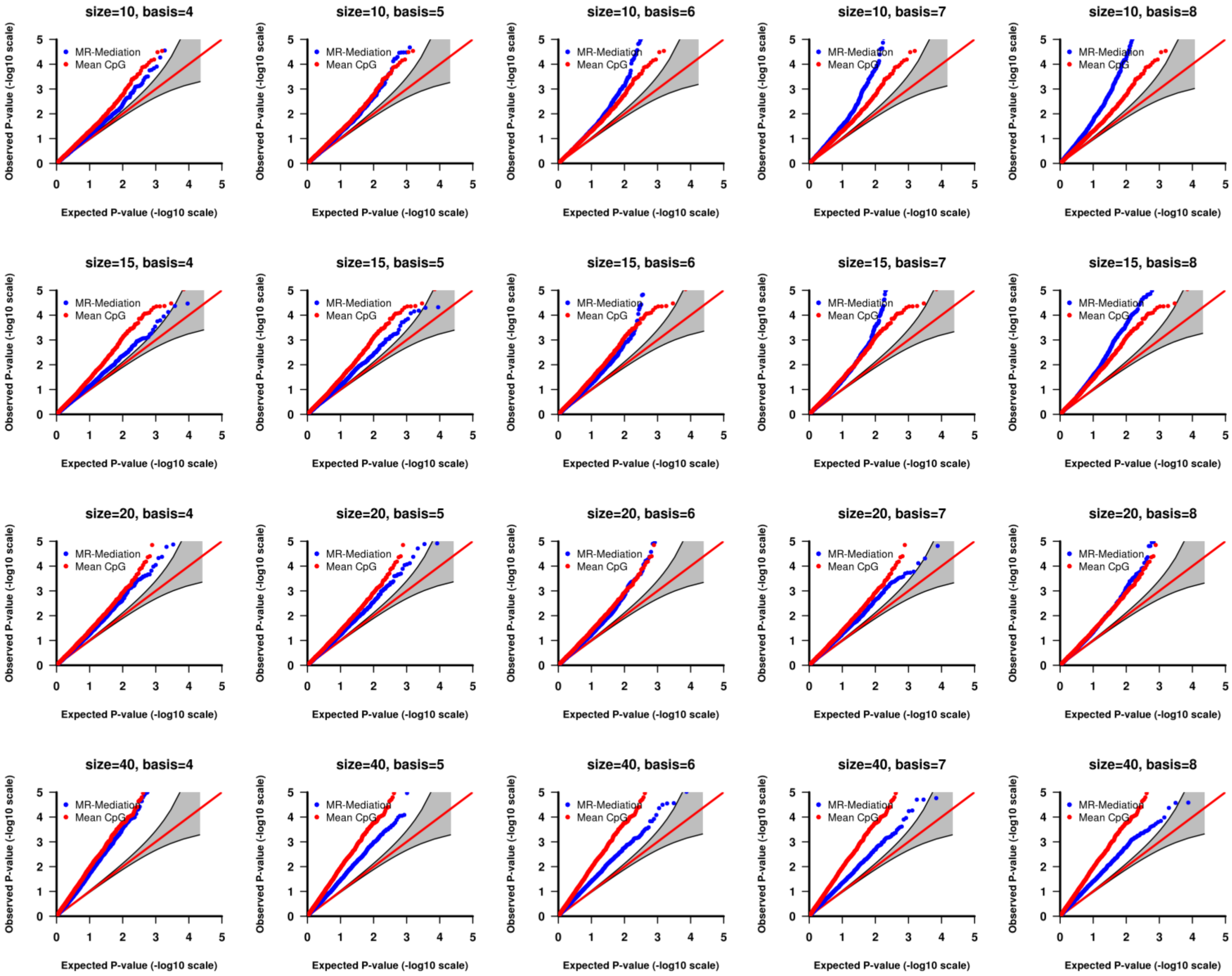


Figure S6. QQ plot of the *p*-values from Setting (2) DE tests with continuous outcome and 100%+ effective CpG sites. Each row shows one gene size with different basis numbers. Two approaches were compared: (1) MR-Mediation; and (2) Mean CpG methylation approach. A 95% pointwise confidence band (gray area) was computed under the assumption that the *p*-values were drawn independently from a uniform [0, 1] distribution.

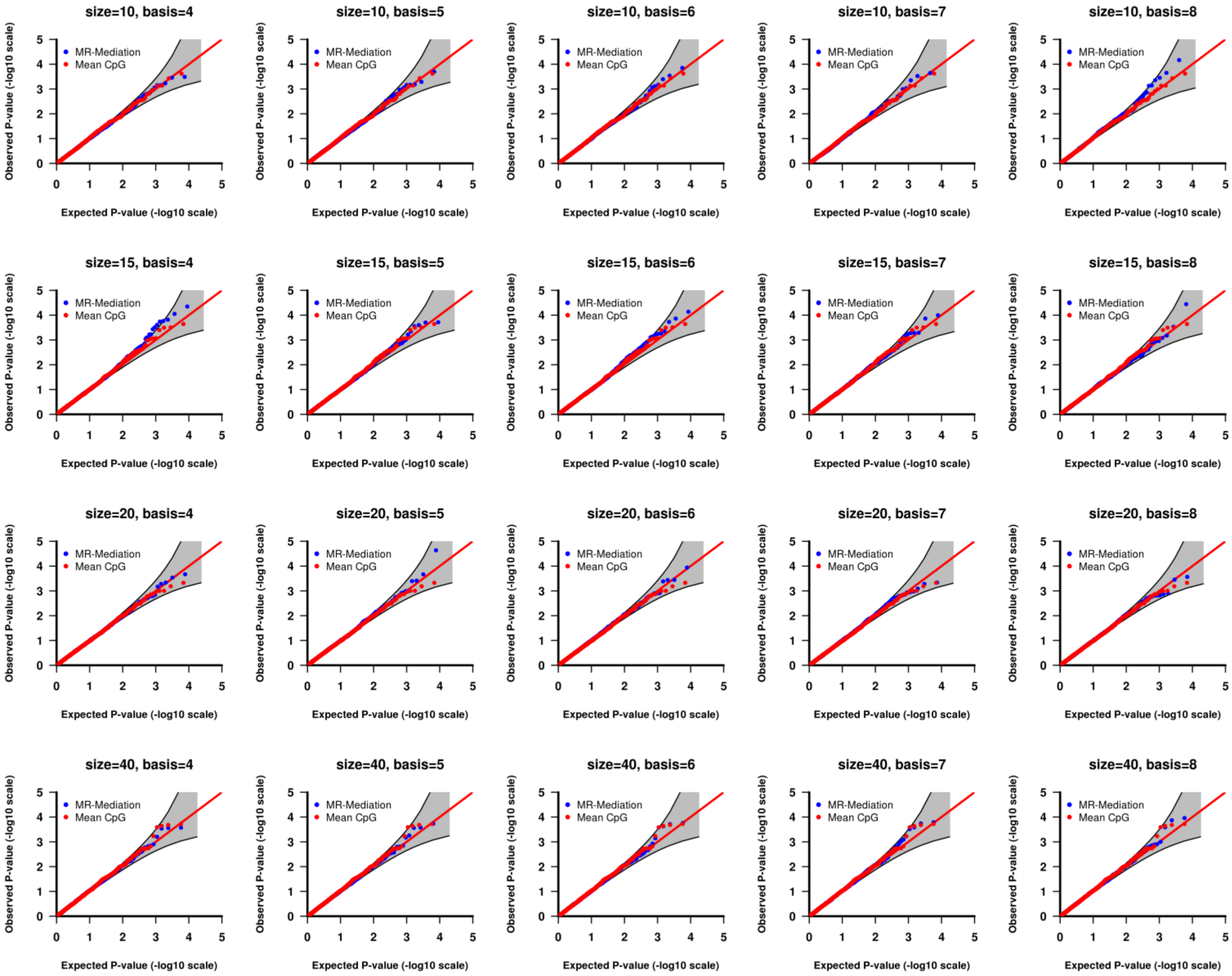


Figure S7. QQ plot of the *p*-values from Setting (3) IE tests with continuous outcome and 100%+ effective CpG sites. Each row shows one gene size with different basis numbers. Two approaches were compared: (1) MR-Mediation; and (2) Mean CpG methylation approach. A 95% pointwise confidence band (gray area) was computed under the assumption that the *p*-values were drawn independently from a uniform [0, 1] distribution.

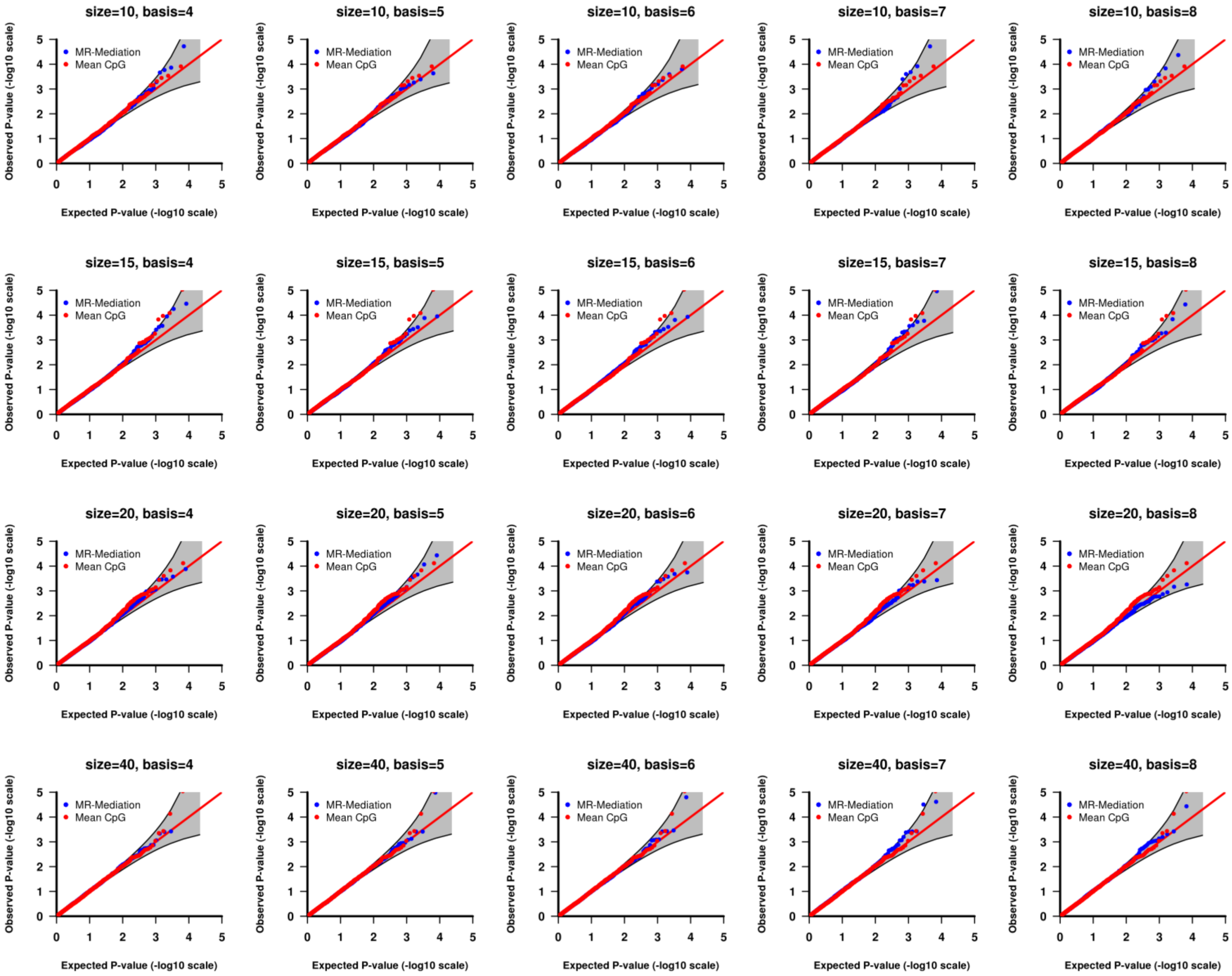


Figure S8. QQ plot of the *p*-values from Setting (1) DE tests with binary outcome. Each row shows one gene size with different basis numbers. Two approaches were compared: (1) MR-Mediation; and (2) Mean CpG methylation approach. A 95% pointwise confidence band (gray area) was computed under the assumption that the *p*-values were drawn independently from a uniform [0, 1] distribution.

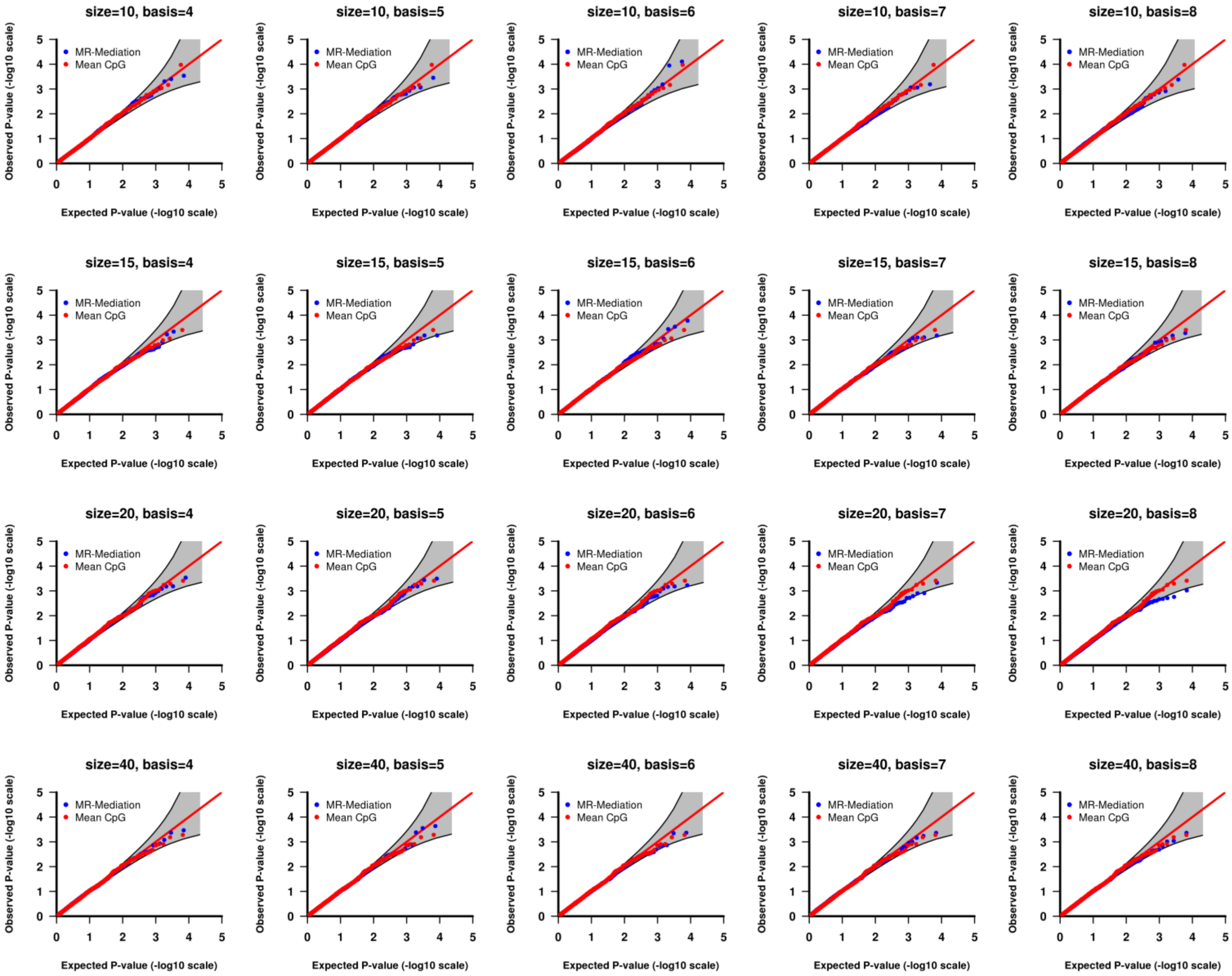


Figure S9. QQ plot of the *p*-values from Setting (1) IE tests with binary outcome. Each row shows one gene size with different basis numbers. Two approaches were compared: (1) MR-Mediation; and (2) Mean CpG methylation approach. A 95% pointwise confidence band (gray area) was computed under the assumption that the *p*-values were drawn independently from a uniform [0, 1] distribution.

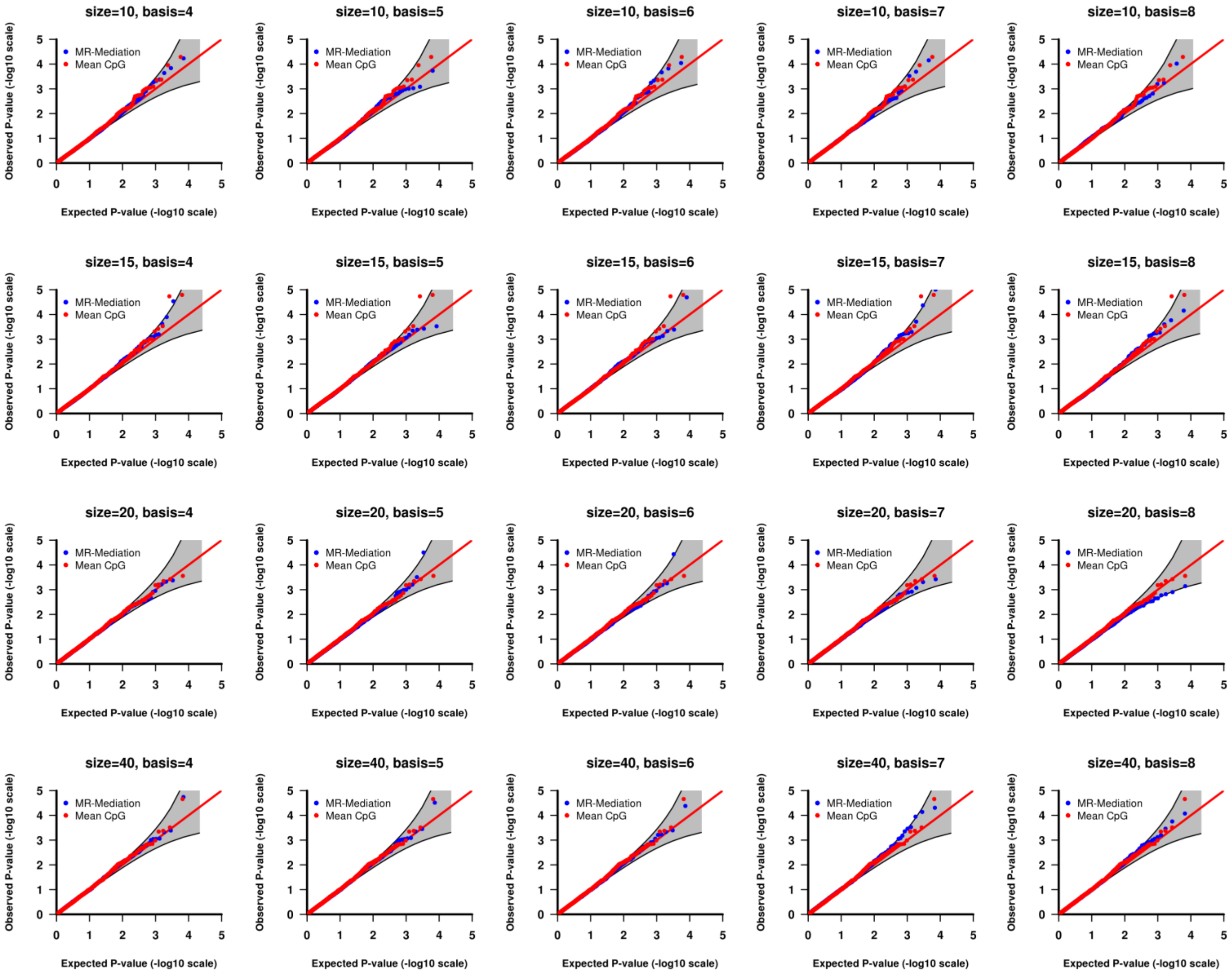


Figure S10. QQ plot of the *p*-values from Setting (1) TE tests with binary outcome. Each row shows one gene size with different basis numbers. Two approaches were compared: (1) MR-Mediation; and (2) Mean CpG methylation approach. A 95% pointwise confidence band (gray area) was computed under the assumption that the *p*-values were drawn independently from a uniform [0, 1] distribution.

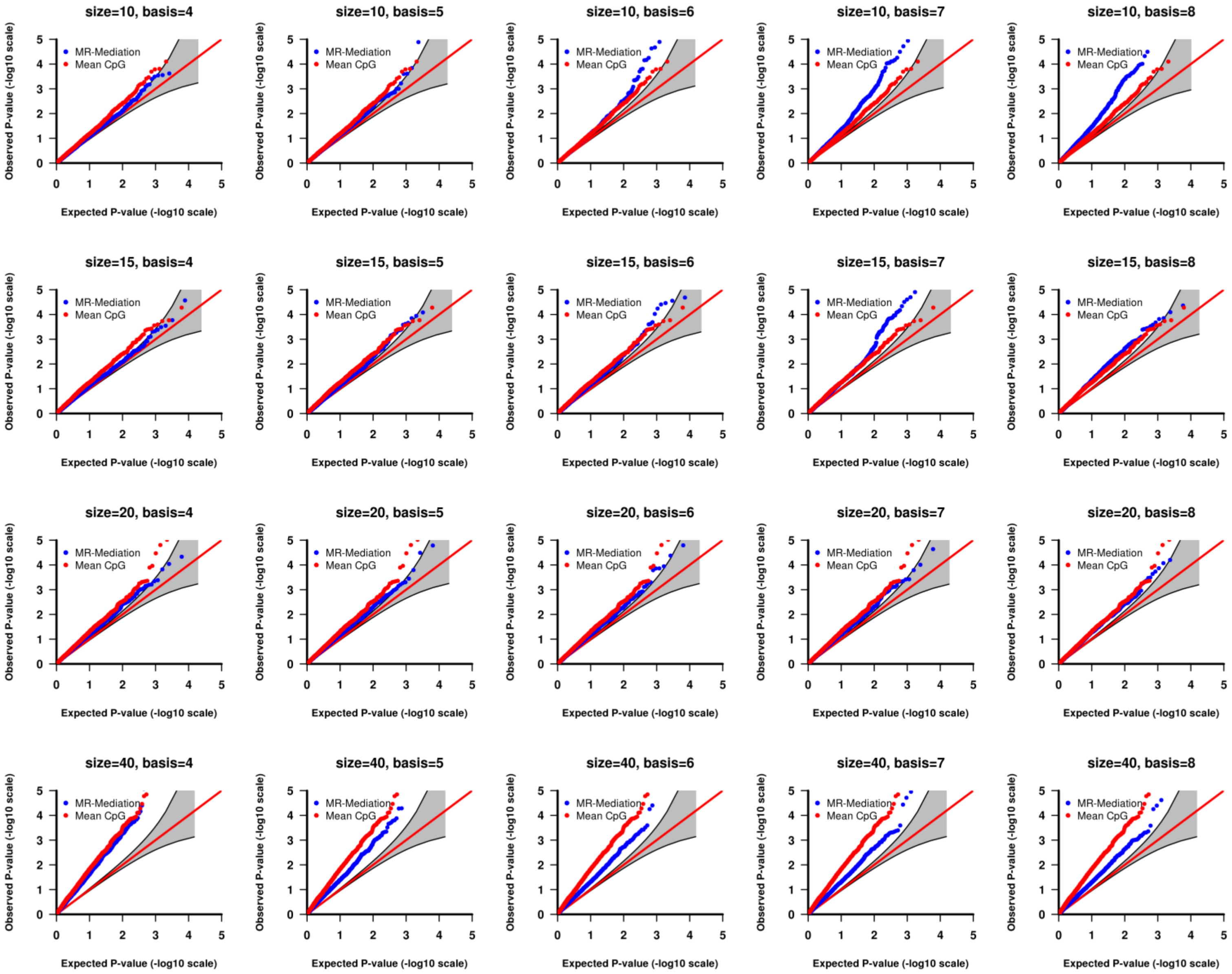


Figure S11. QQ plot of the *p*-values from Setting (2) DE tests with binary outcome and 100%+ effective CpG sites. Each row shows one gene size with different basis numbers. Two approaches were compared: (1) MR-Mediation; and (2) Mean CpG methylation approach. A 95% pointwise confidence band (gray area) was computed under the assumption that the *p*-values were drawn independently from a uniform [0, 1] distribution.

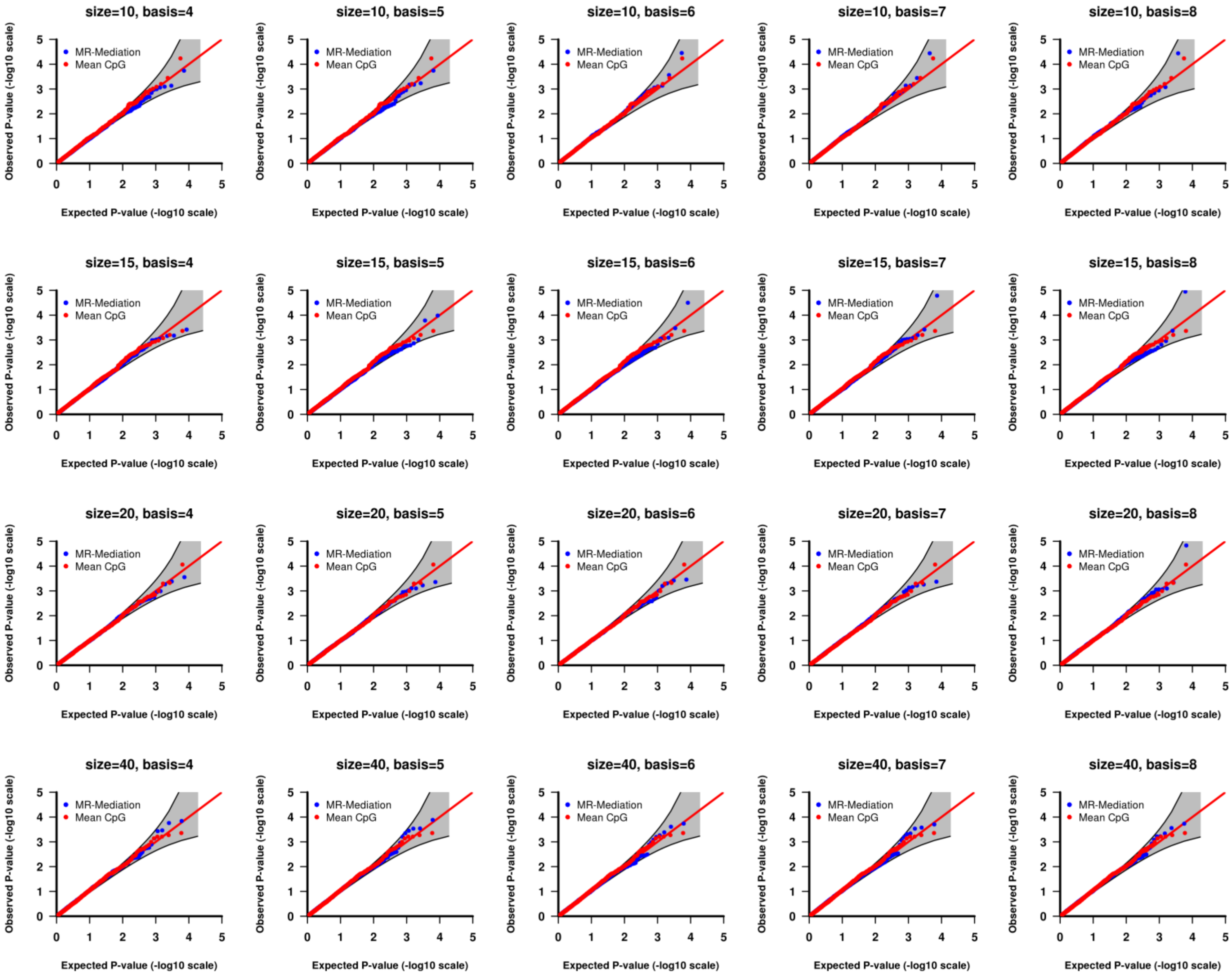


Figure S12. QQ plot of the *p*-values from Setting (3) IE tests with binary outcome and 100%+ effective CpG sites. Each row shows one gene size with different basis numbers. Two approaches were compared: (1) MR-Mediation; and (2) Mean CpG methylation approach. A 95% pointwise confidence band (gray area) was computed under the assumption that the *p*-values were drawn independently from a uniform [0, 1] distribution.

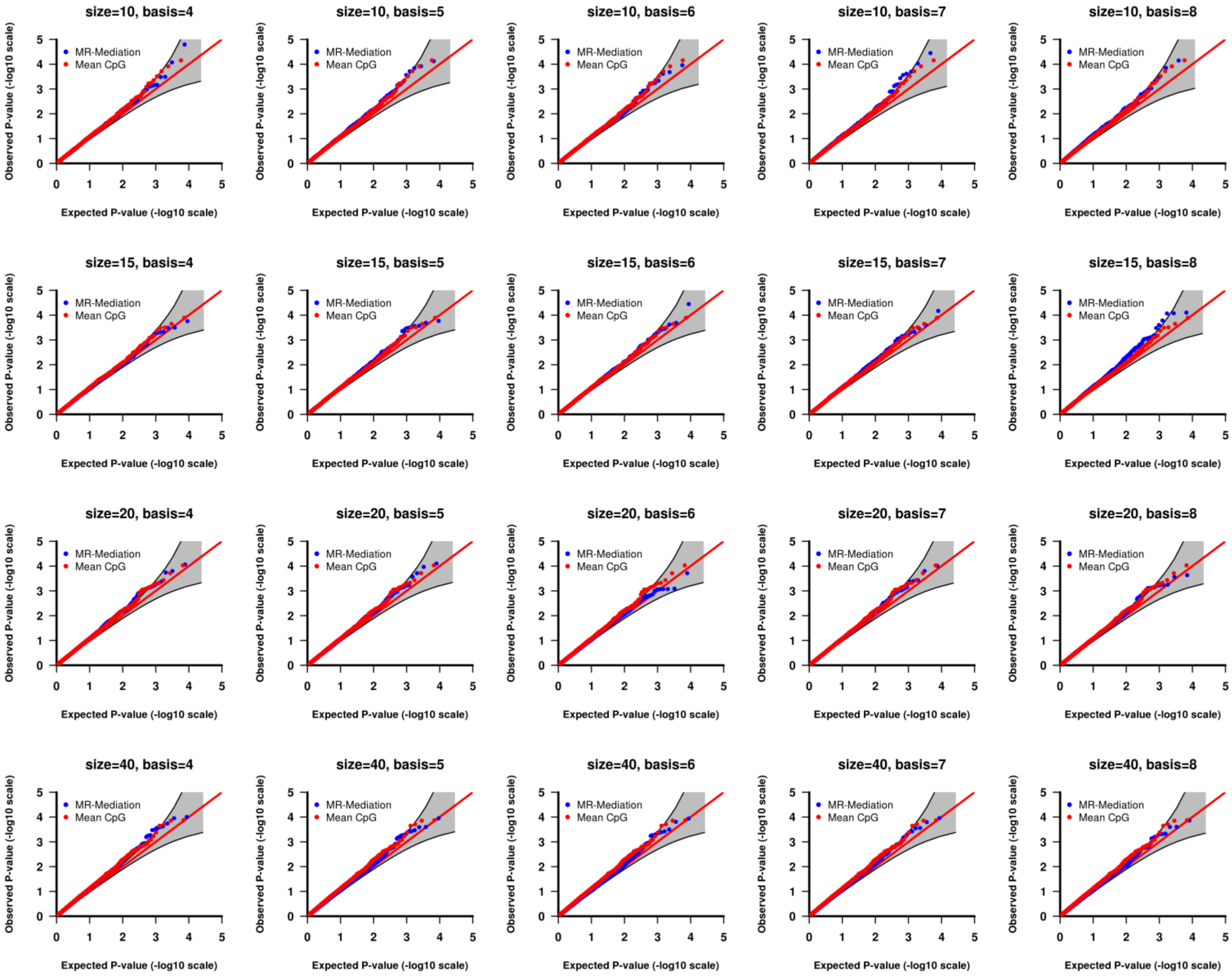


Figure S13. QQ plot of the *p*-values from Setting (2) DE tests with continuous outcome and 50%+/50%- effective CpG sites. Each row shows one gene size with different basis numbers. Two approaches were compared: (1) MR-Mediation; and (2) Mean CpG methylation approach. A 95% pointwise confidence band (gray area) was computed under the assumption that the *p*-values were drawn independently from a uniform [0, 1] distribution.

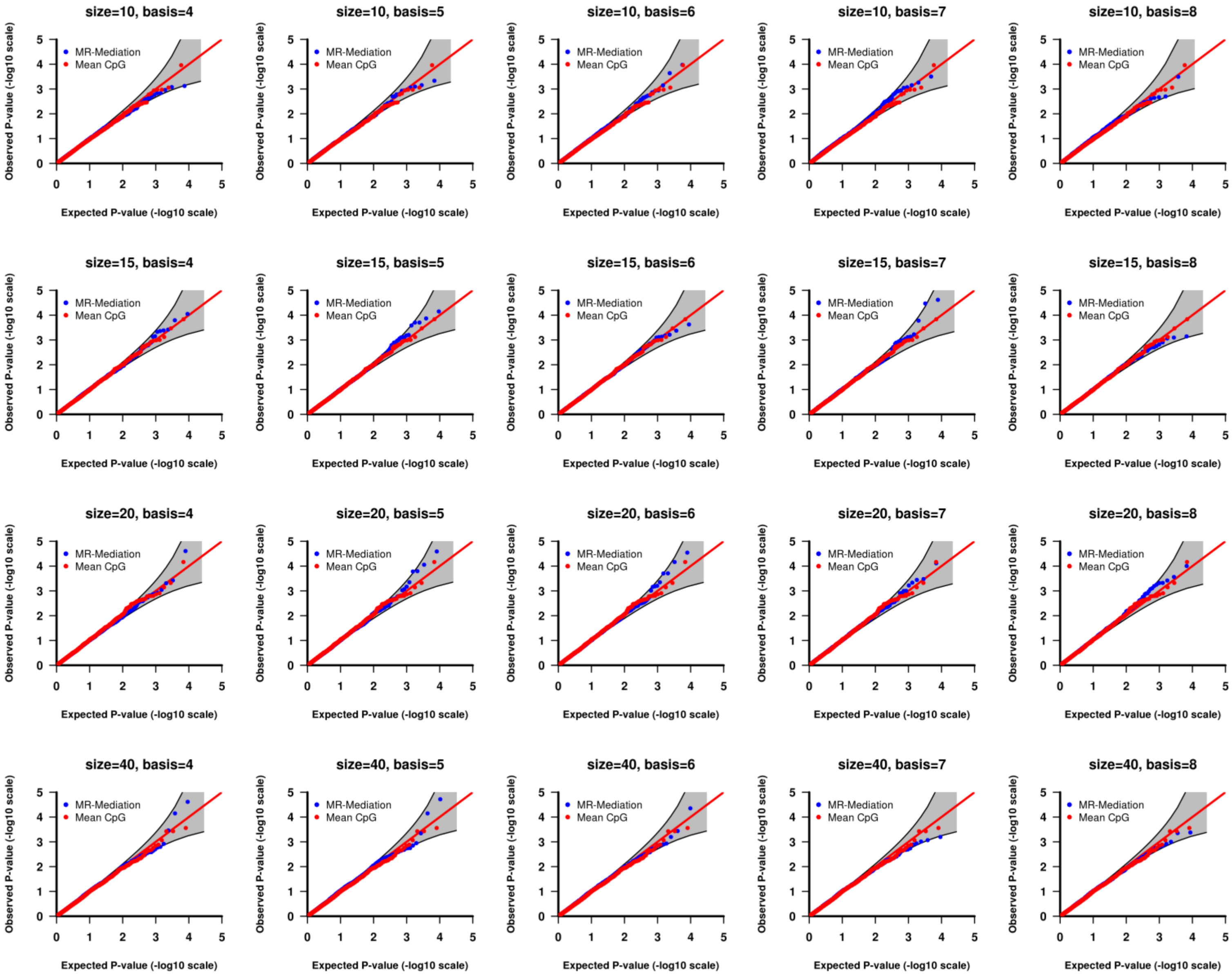


Figure S14. QQ plot of the *p*-values from Setting (3) IE tests with continuous outcome and 50%+/50%- effective CpG sites. Each row shows one gene size with different basis numbers. Two approaches were compared: (1) MR-Mediation; and (2) Mean CpG methylation approach. A 95% pointwise confidence band (gray area) was computed under the assumption that the *p*-values were drawn independently from a uniform [0, 1] distribution.

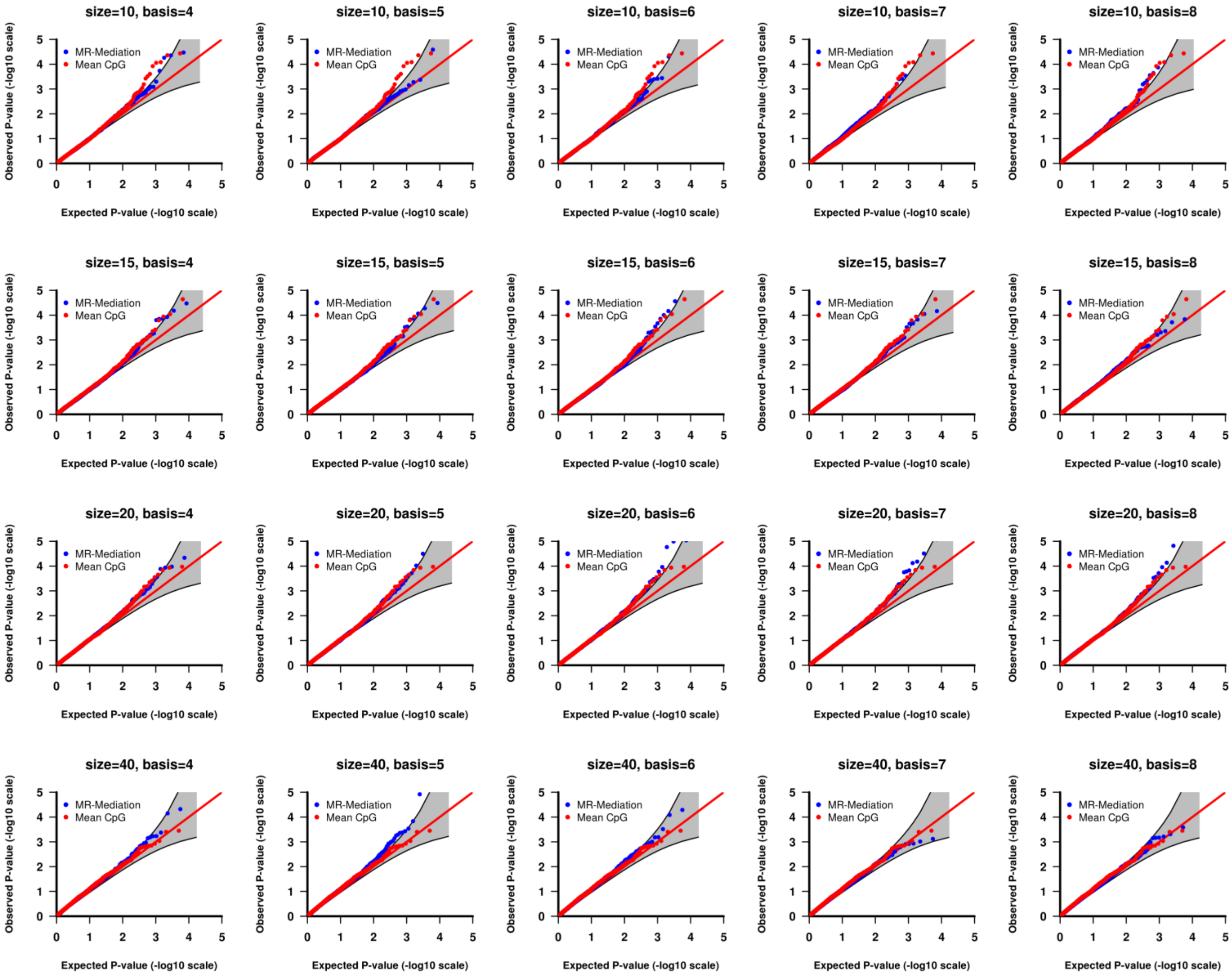


Figure S15. QQ plot of the *p*-values from Setting (2) DE tests with binary outcome and 50%+/50%- effective CpG sites. Each row shows one gene size with different basis numbers. Two approaches were compared: (1) MR-Mediation; and (2) Mean CpG methylation approach. A 95% pointwise confidence band (gray area) was computed under the assumption that the *p*-values were drawn independently from a uniform [0, 1] distribution.

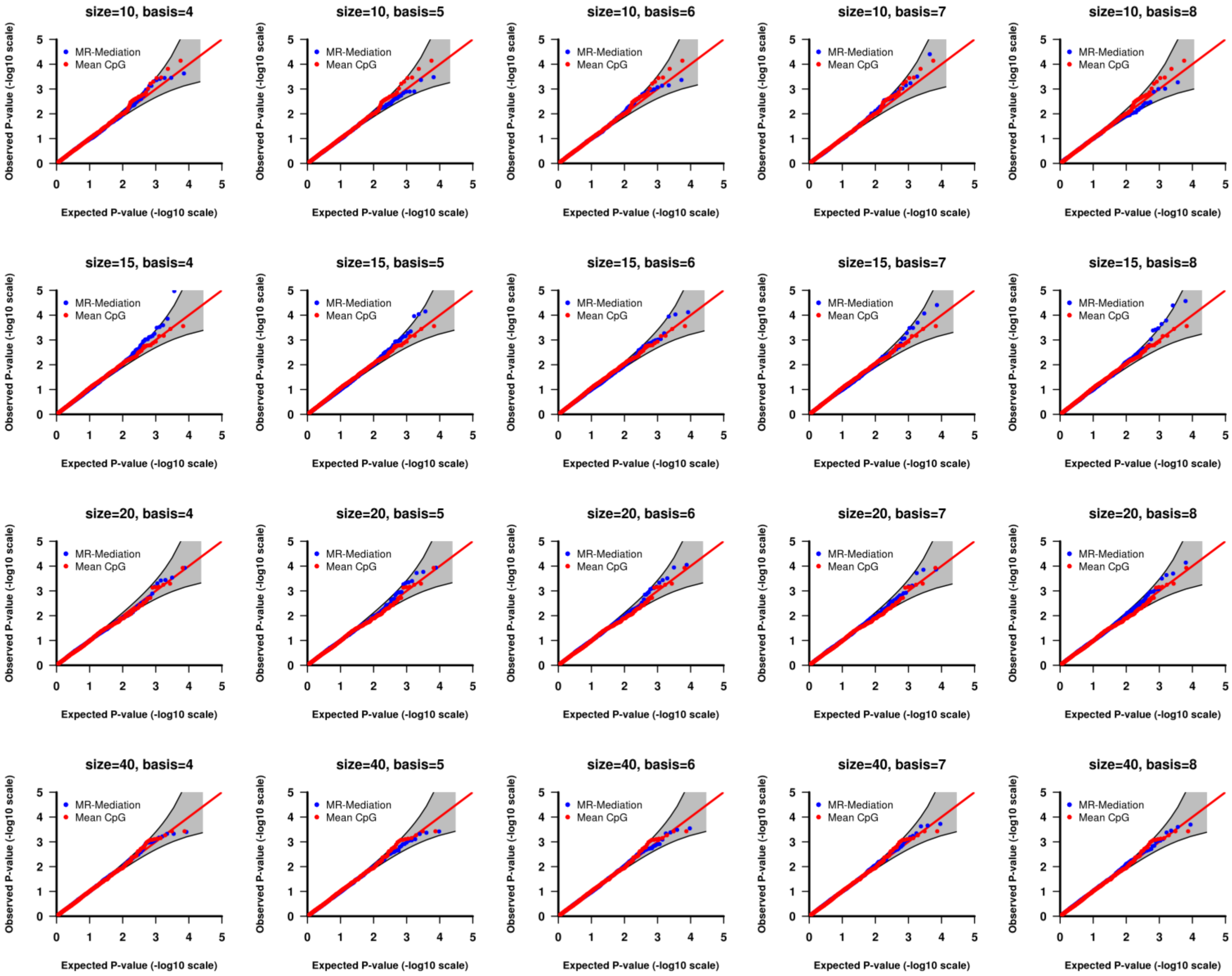


Figure S16. QQ plot of the *p*-values from Setting (3) IE tests with binary outcome and 50%+/50%- effective CpG sites. Each row shows one gene size with different basis numbers. Two approaches were compared: (1) MR-Mediation; and (2) Mean CpG methylation approach. A 95% pointwise confidence band (gray area) was computed under the assumption that the *p*-values were drawn independently from a uniform [0, 1] distribution.

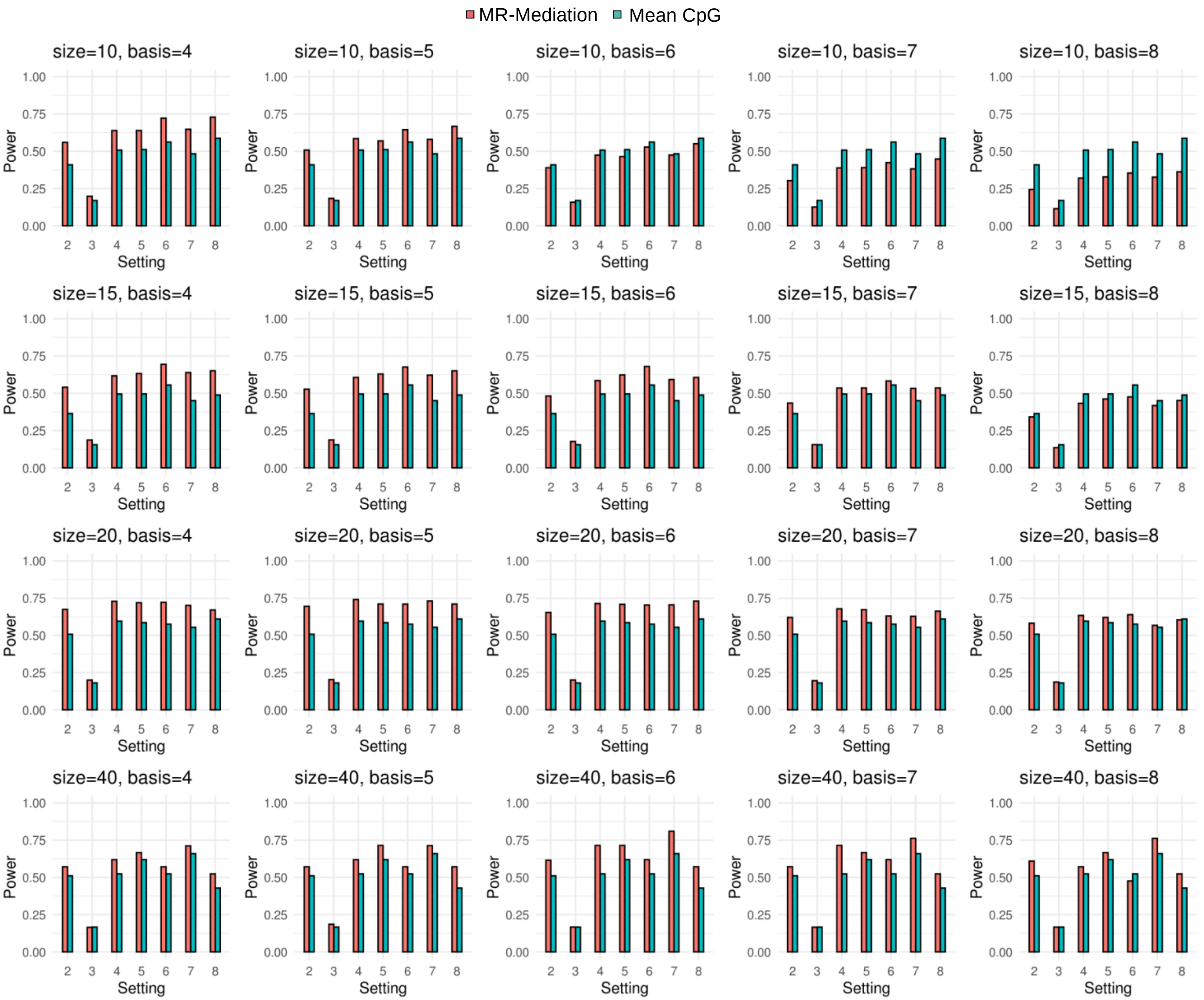


Figure S17. Power comparison for TE with continuous outcome and 100%+ effective CpG sites. Each row shows one gene size with different basis numbers. Two approaches were compared: (1) MR-Mediation; and (2) Mean CpG methylation approach.

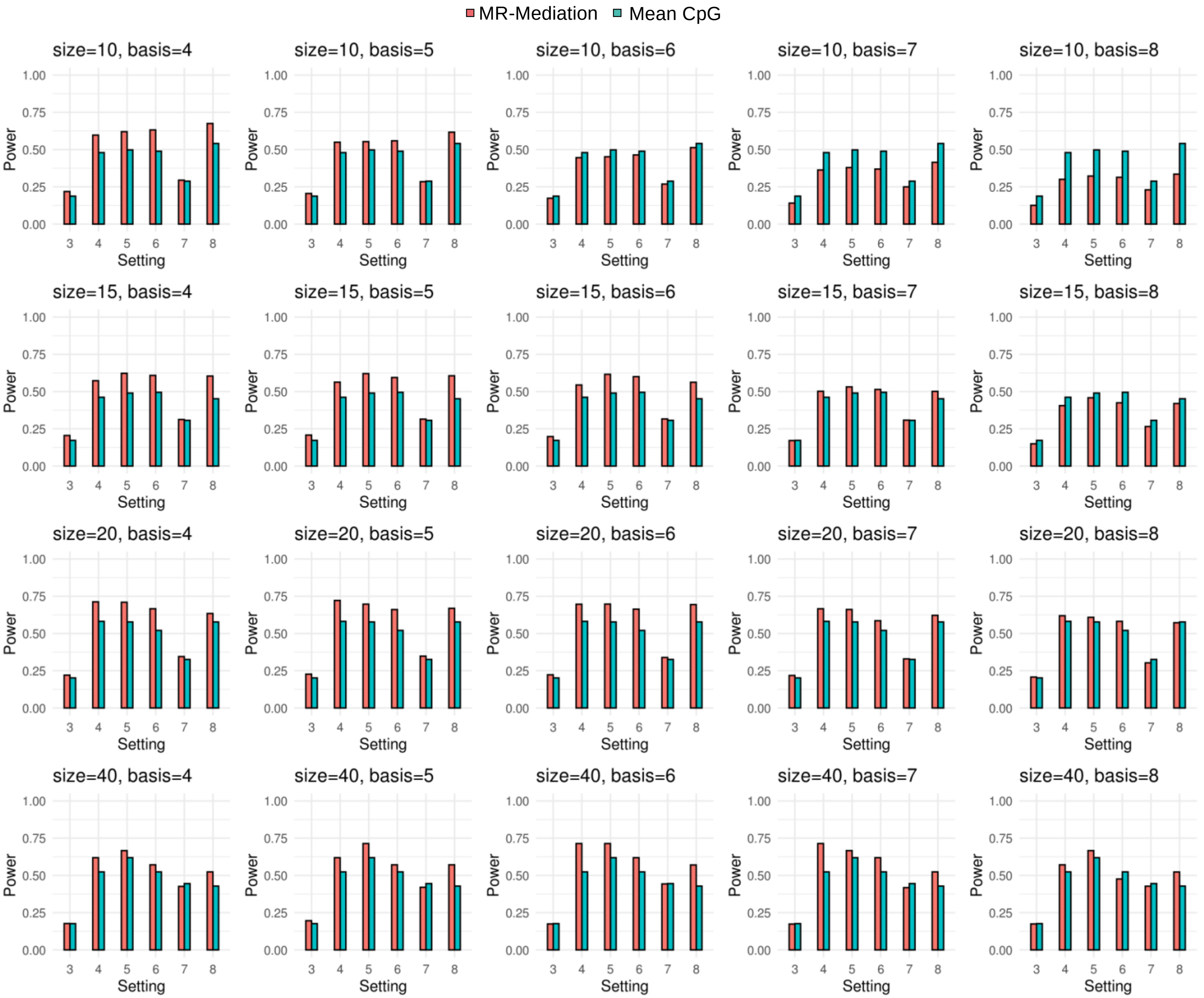


Figure S18. Power comparison for DE with continuous outcome and 100%+ effective CpG sites. Each row shows one gene size with different basis numbers. Two approaches were compared: (1) MR-Mediation; and (2) Mean CpG methylation approach.

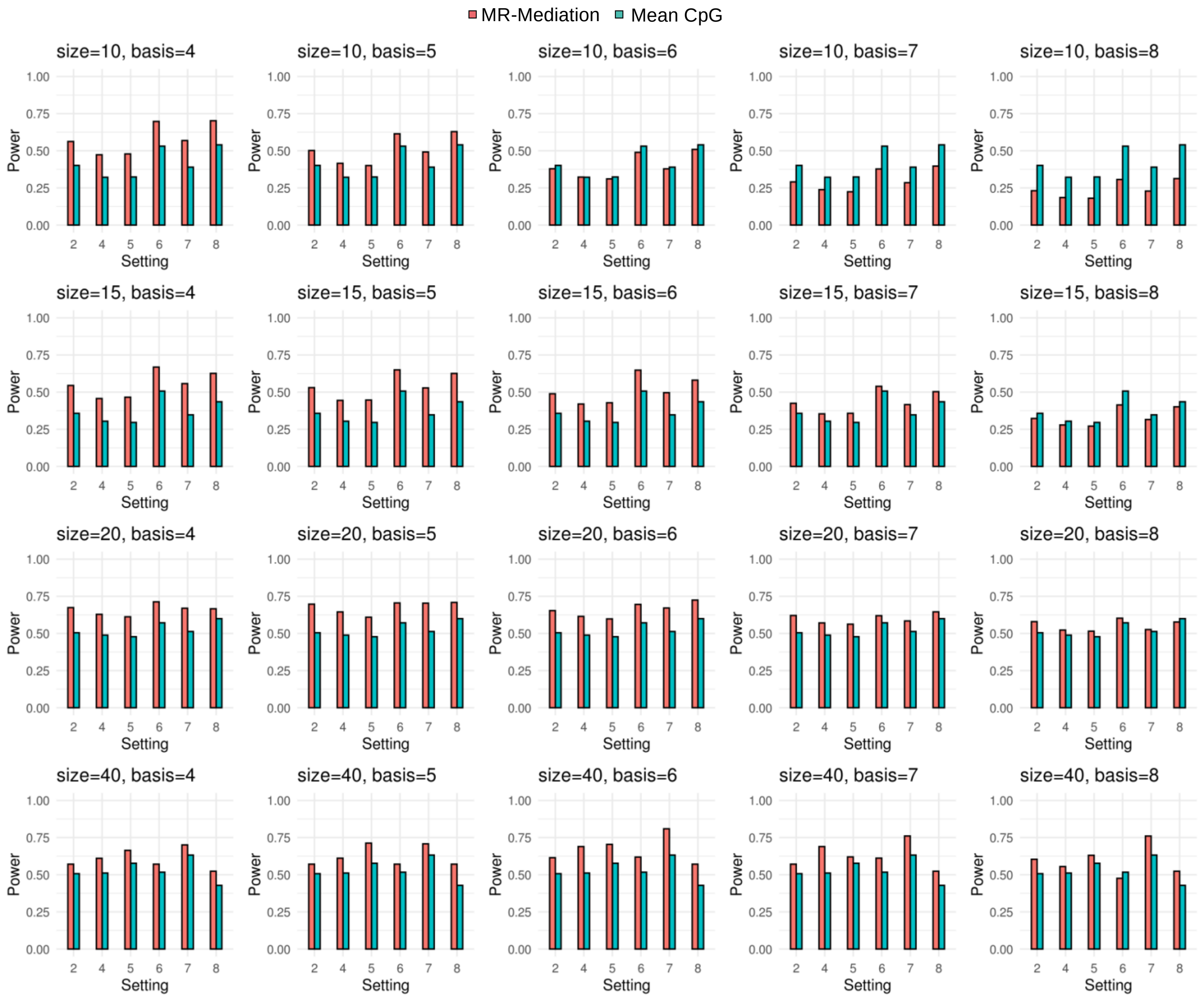


Figure S19. Power comparison for IE with continuous outcome and 100%+ effective CpG sites. Each row shows one gene size with different basis numbers. Two approaches were compared: (1) MR-Mediation; and (2) Mean CpG methylation approach.

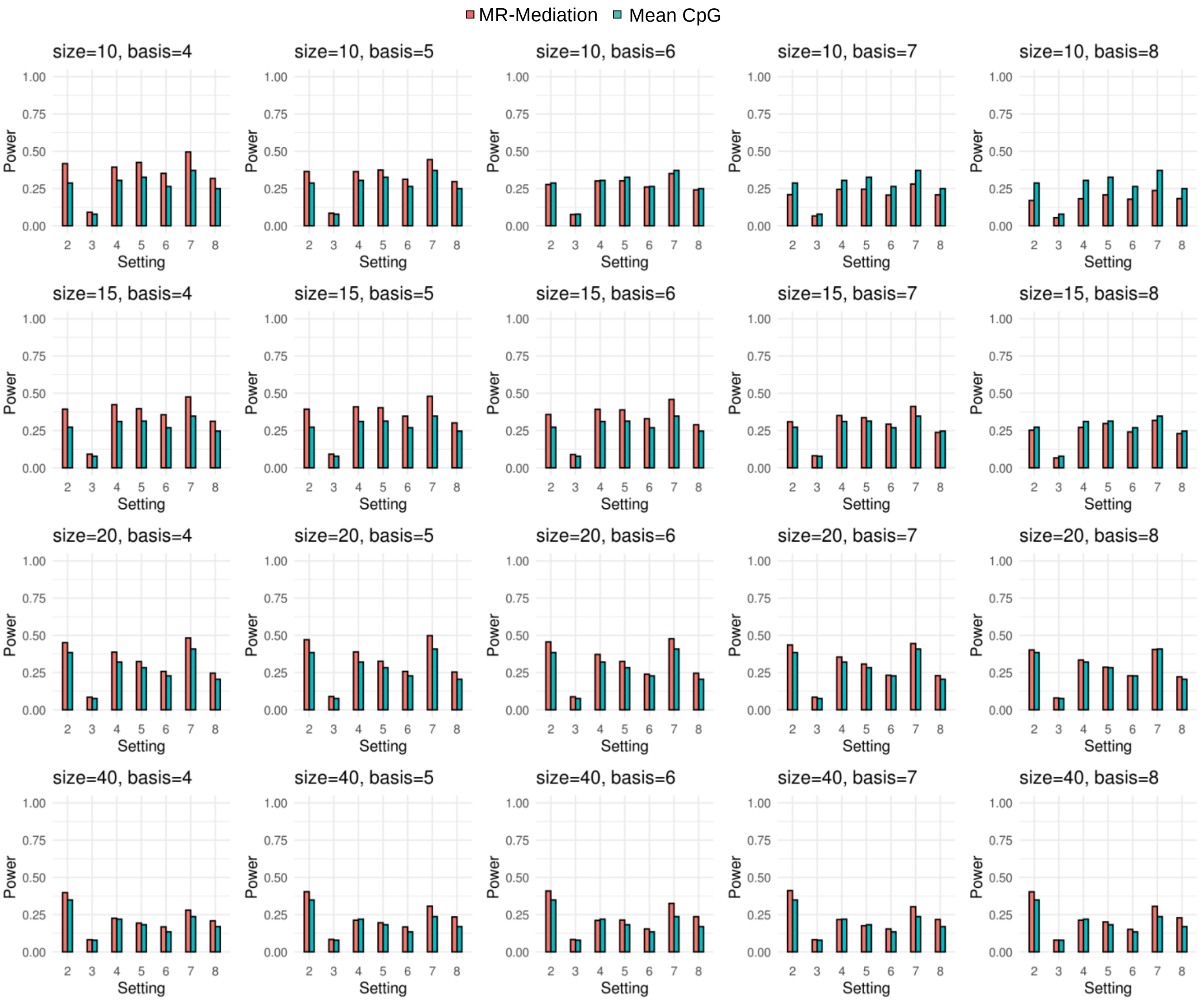


Figure S20. Power comparison for TE with binary outcome and 100%+ effective CpG sites. Each row shows one gene size with different basis numbers. Two approaches were compared: (1) MR-Mediation; and (2) Mean CpG methylation approach.

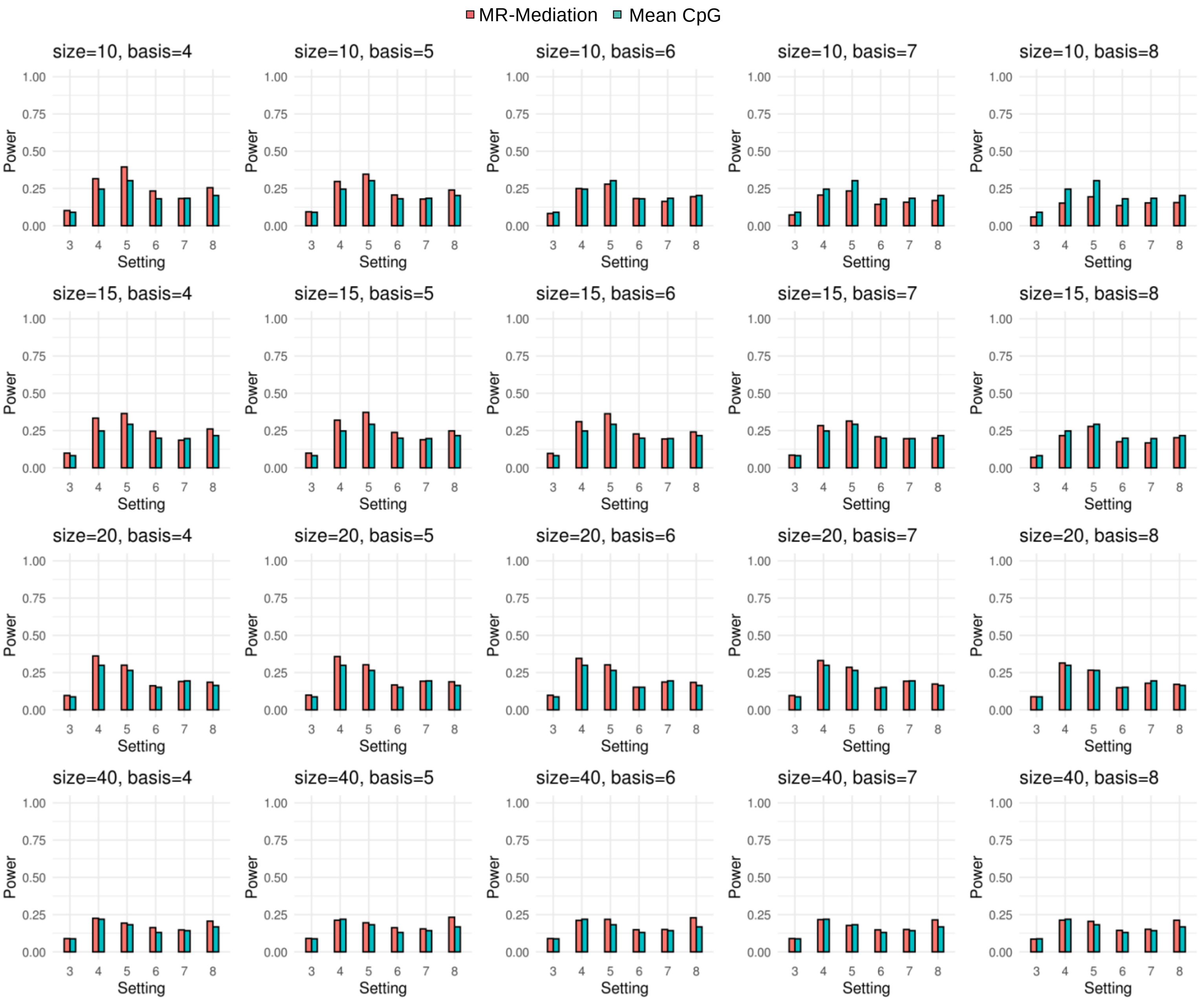


Figure S21. Power comparison for DE with binary outcome and 100%+ effective CpG sites. Each row shows one gene size with different basis numbers. Two approaches were compared: (1) MR-Mediation; and (2) Mean CpG methylation approach.

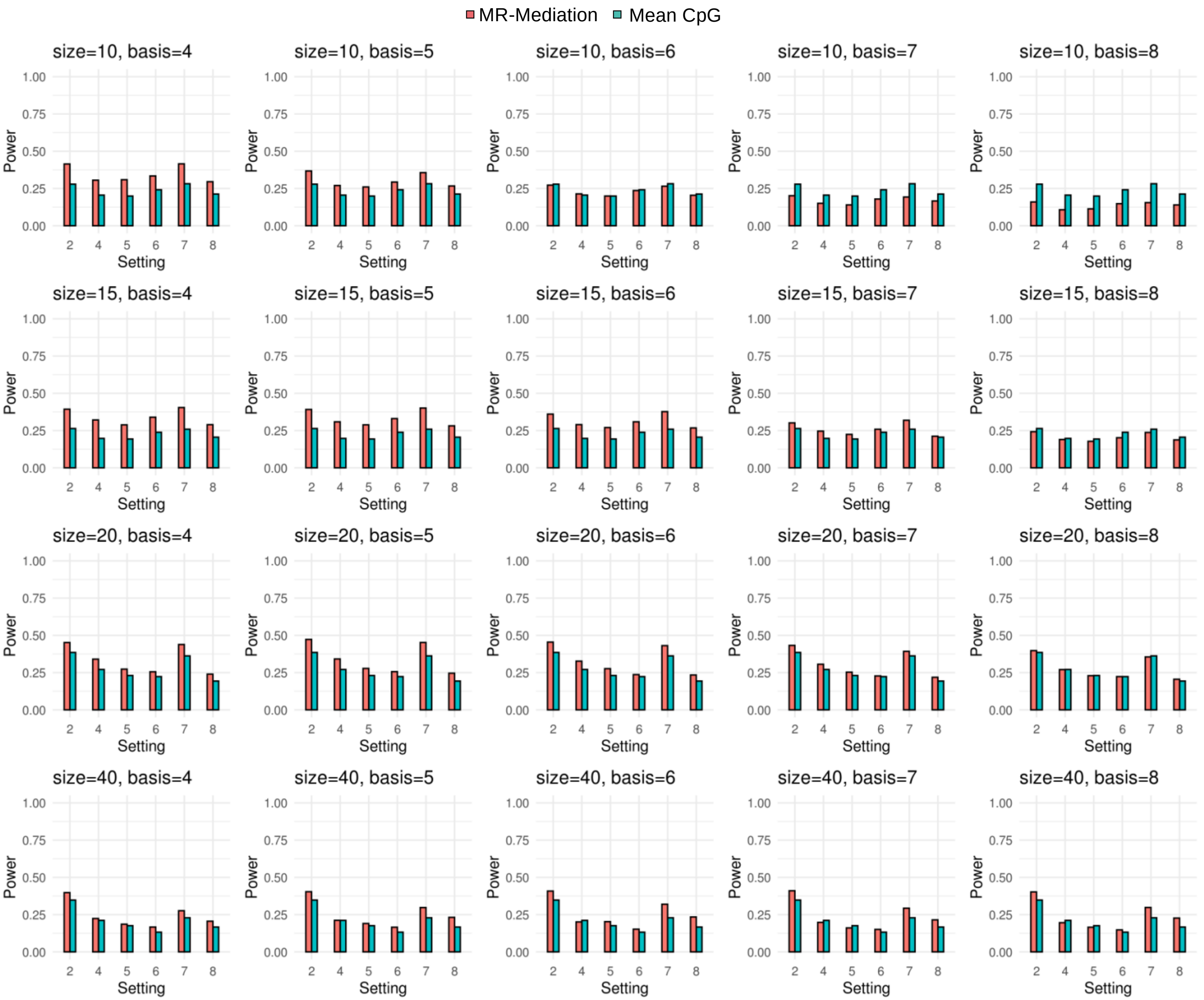


Figure S22. Power comparison for IE with binary outcome and 100%+ effective CpG sites. Each row shows one gene size with different basis numbers. Two approaches were compared: (1) MR-Mediation; and (2) Mean CpG methylation approach.

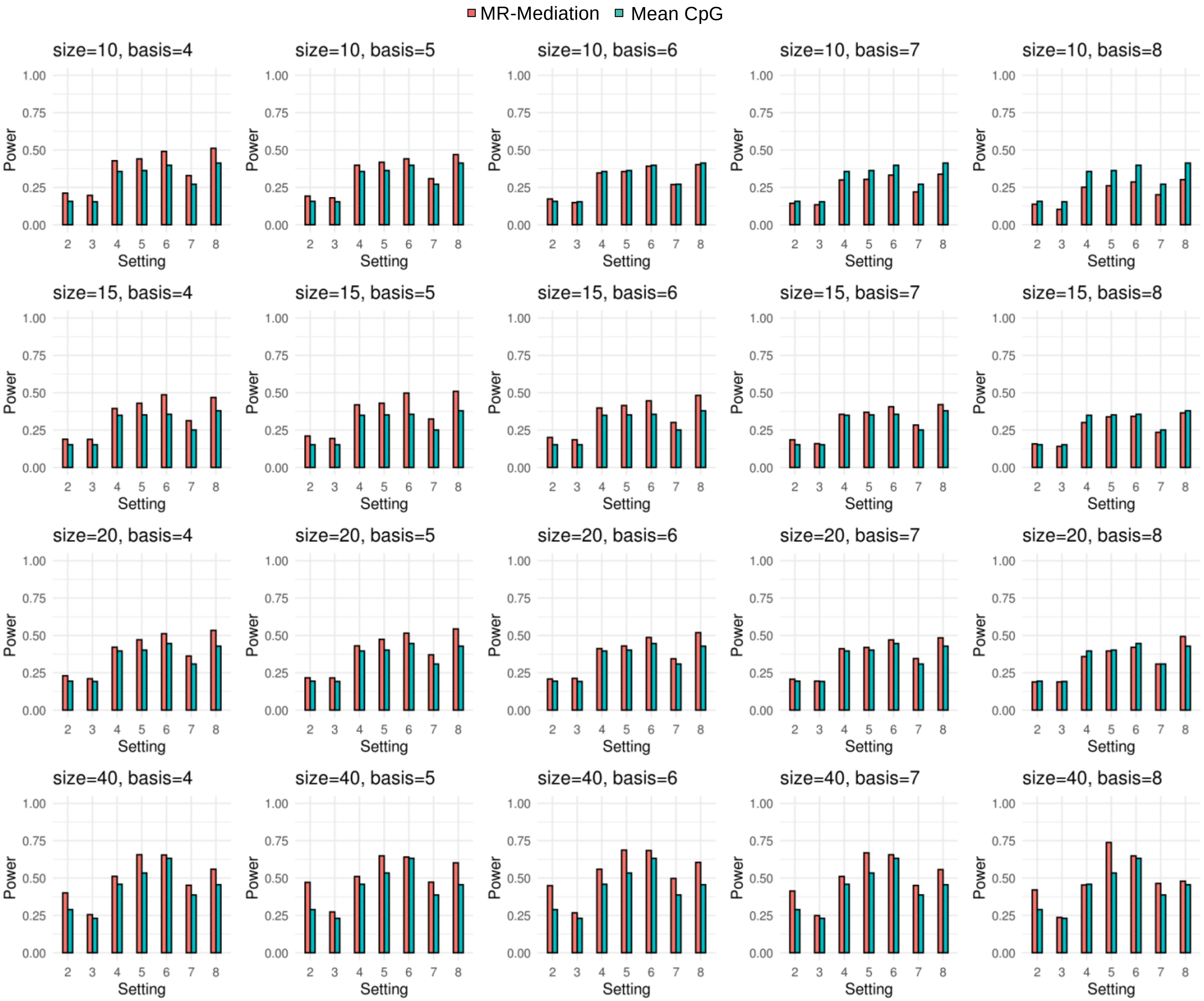


Figure S23. Power comparison for TE with continuous outcome and 50%+/50%- effective CpG sites. Each row shows one gene size with different basis numbers. Two approaches were compared: (1) MR-Mediation; and (2) Mean CpG methylation approach.

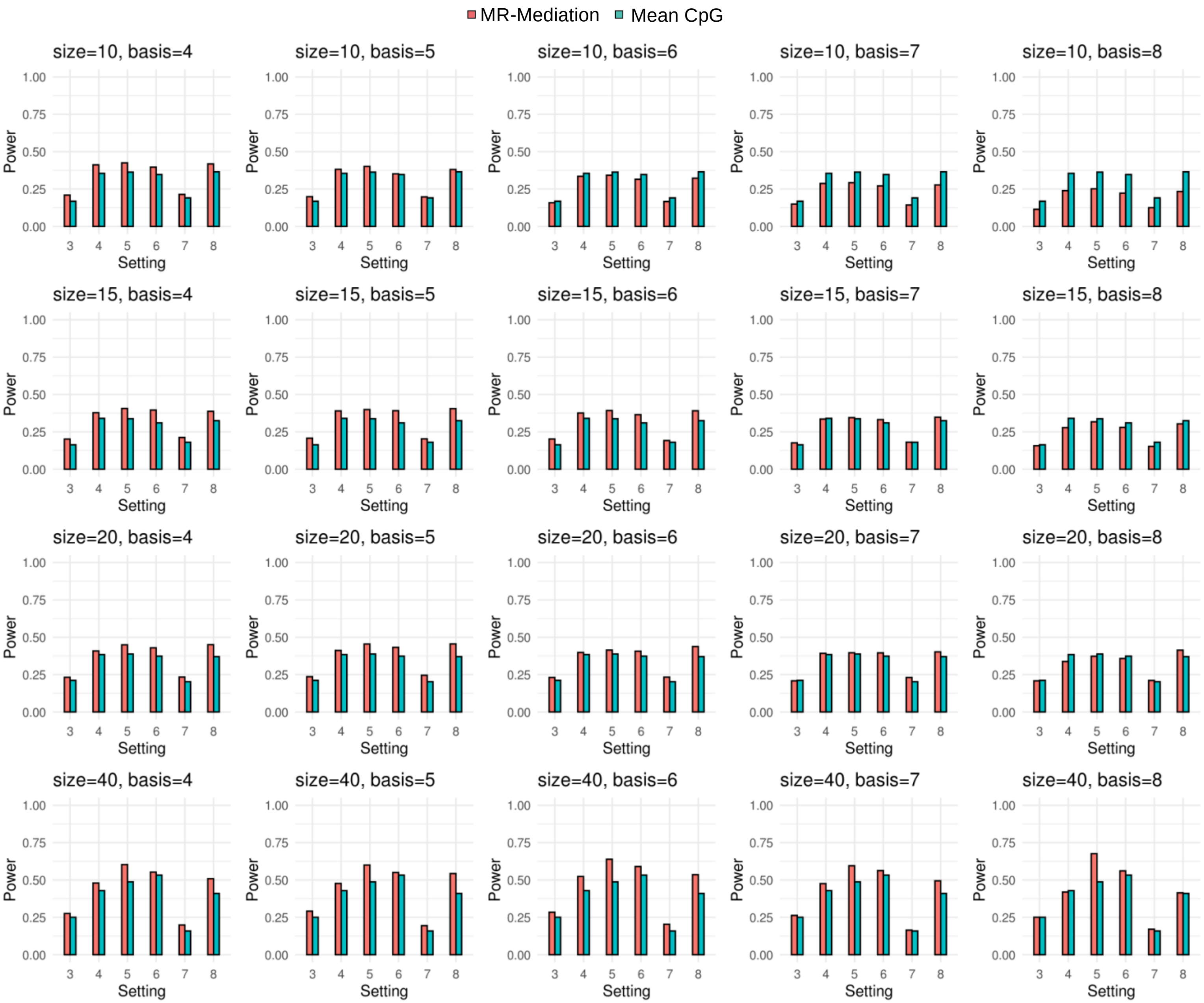


Figure S24. Power comparison for DE with continuous outcome and 50%+/50%- effective CpG sites. Each row shows one gene size with different basis numbers. Two approaches were compared: (1) MR-Mediation; and (2) Mean CpG methylation approach.

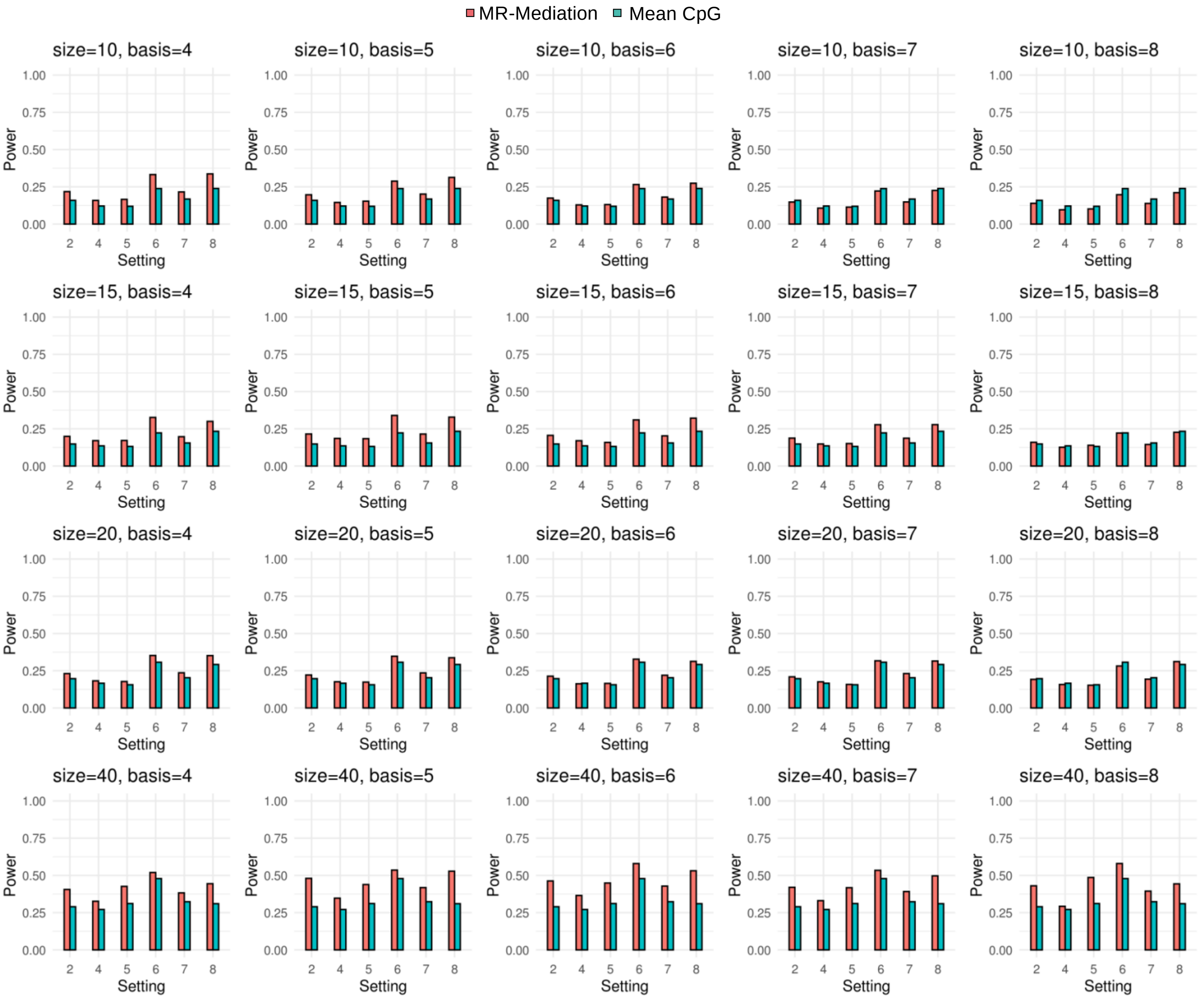


Figure S25. Power comparison for IE with continuous outcome and 50%+/50%- effective CpG sites. Each row shows one gene size with different basis numbers. Two approaches were compared: (1) MR-Mediation; and (2) Mean CpG methylation approach.

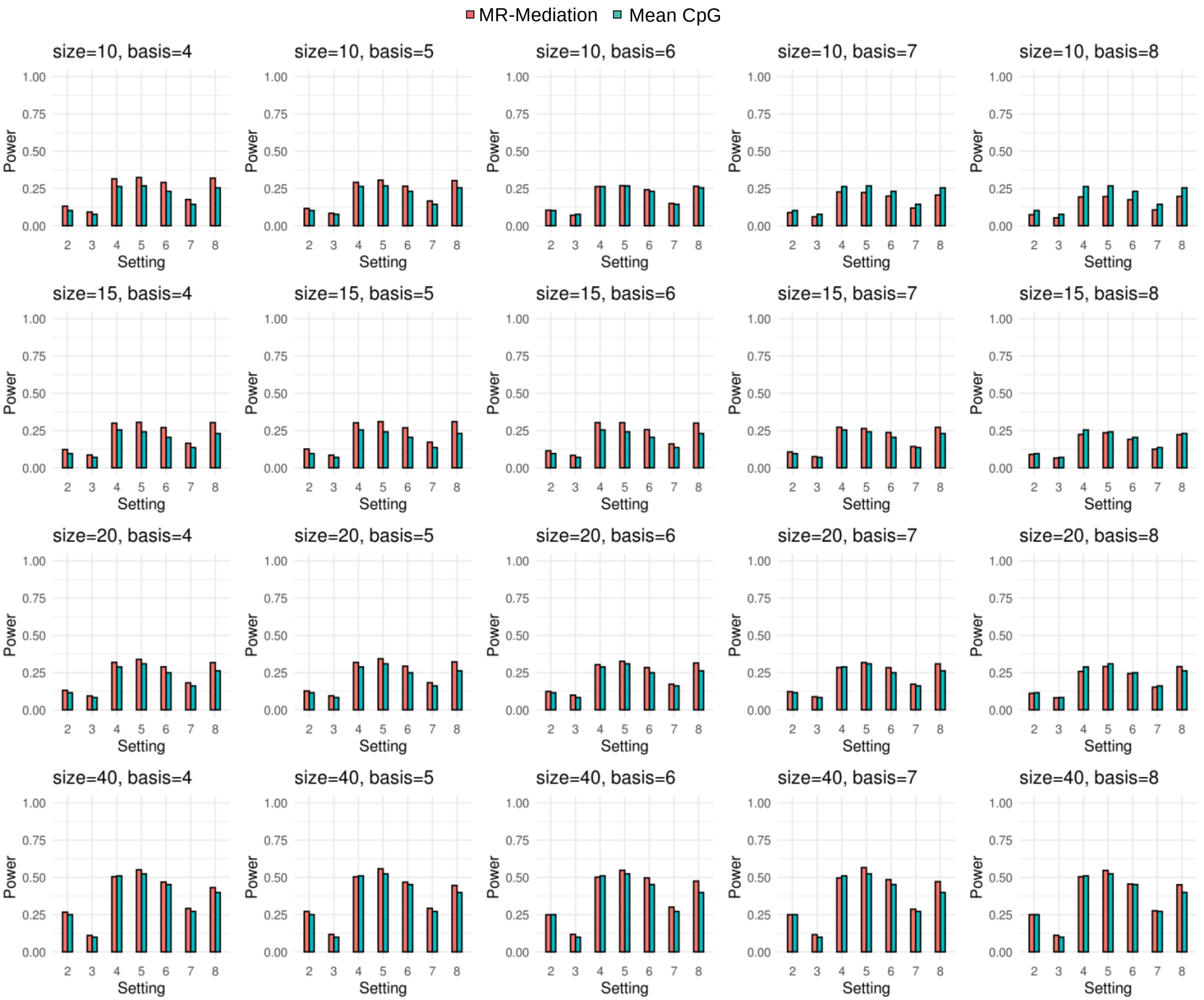


Figure S26. Power comparison for TE with binary outcome and 50%+/50%- effective CpG sites. Each row shows one gene size with different basis numbers. Two approaches were compared: (1) MR-Mediation; and (2) Mean CpG methylation approach.

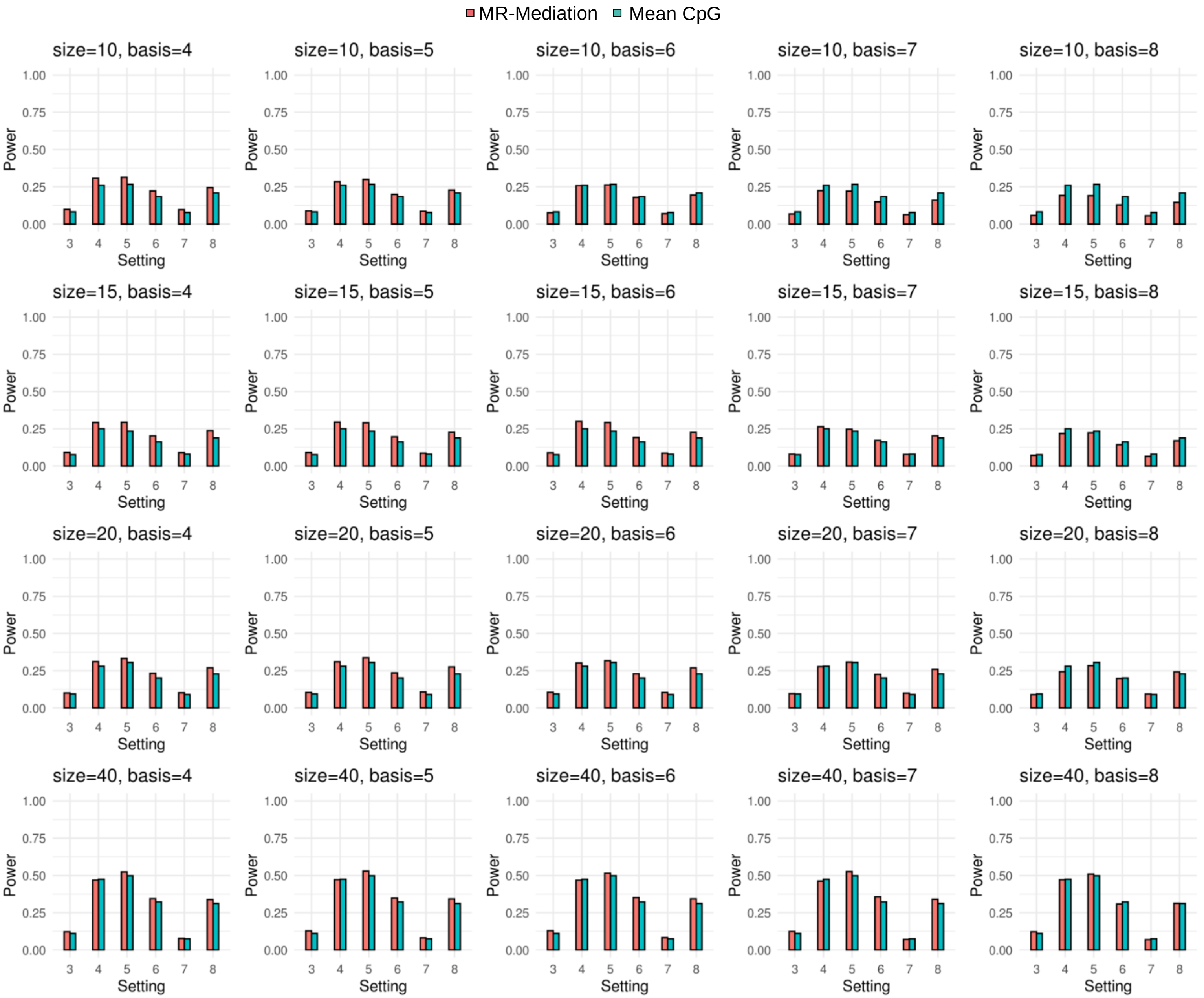


Figure S27. Power comparison for DE with binary outcome and 50%+/50%- effective CpG sites. Each row shows one gene size with different basis numbers. Two approaches were compared: (1) MR-Mediation; and (2) Mean CpG methylation approach.

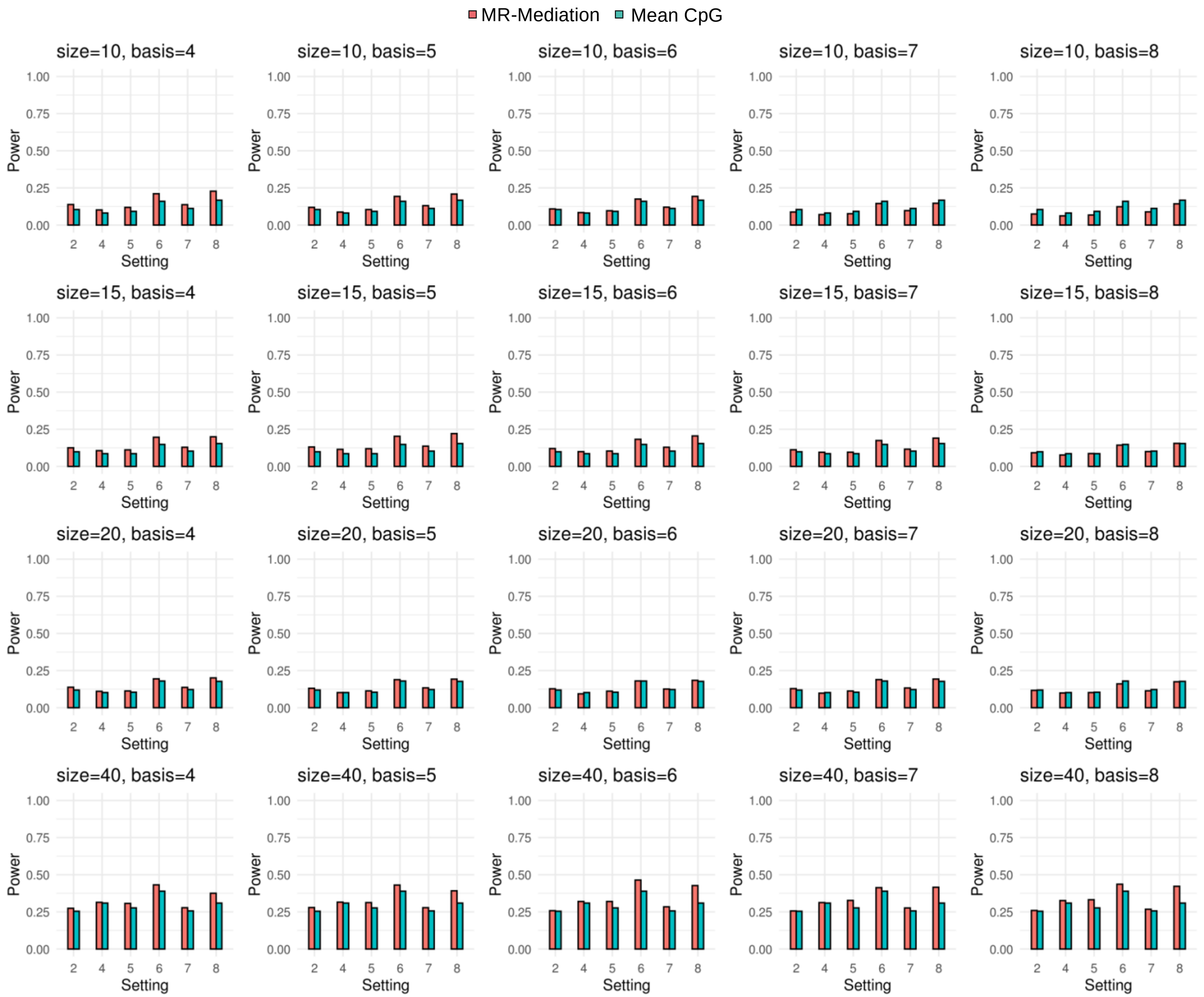


Figure S28. Power comparison for IE with binary outcome and 50%+/50%- effective CpG sites. Each row shows one gene size with different basis numbers. Two approaches were compared: (1) MR-Mediation; and (2) Mean CpG methylation approach.

Table S1. The gene-based methylation mediated effect (IE) on the effect of exposure to gun violence on atopic asthma (i.e., gun → methylation → AA) from top 20 association pairs between exposure to gun violence and gene-based methylation in non-asthmatic subjects (i.e., gun → methylation).

| Gene | Size | Chr | Position | *P*_XM* | *FDR*_XM | *P*_IE |
|---|---|---|---|---|---|---|
| *CFD* | 10 | chr19 | 859330 | 1.93E-05 | 0.1102 | 3.09E-02 |
| *TBC1D14* | 38 | chr4 | 6909491 | 3.53E-04 | 0.9984 | 7.28E-03 |
| *FAM120B* | 13 | chr6 | 170614479 | 9.05E-04 | 0.9984 | 3.33E-02 |
| *LINC00944* | 11 | chr12 | 127212034 | 1.08E-03 | 0.9984 | 4.84E-01 |
| *CRB1* | 10 | chr1 | 197173499 | 1.69E-03 | 0.9984 | 1.75E-02 |
| *FOXI1* | 13 | chr5 | 169529408 | 1.72E-03 | 0.9984 | 1.61E-01 |
| *RP1* | 11 | chr8 | 55527869 | 2.02E-03 | 0.9984 | 5.48E-01 |
| *LOC339166* | 10 | chr17 | 5673550 | 2.10E-03 | 0.9984 | 1.42E-02 |
| *ZGPAT* | 15 | chr20 | 62340198 | 2.29E-03 | 0.9984 | 2.55E-03 |
| *DZIP1* | 10 | chr13 | 96234082 | 2.31E-03 | 0.9984 | 7.83E-01 |
| *MED16* | 11 | chr19 | 865897 | 2.44E-03 | 0.9984 | 2.05E-02 |
| *TBPL2* | 13 | chr14 | 55885957 | 3.21E-03 | 0.9984 | 1.50E-01 |
| *MIR143HG* | 20 | chr5 | 148785250 | 3.70E-03 | 0.9984 | 1.29E-01 |
| *PCDHB2* | 10 | chr5 | 140472925 | 4.63E-03 | 0.9984 | 1.44E-01 |
| *PLEKHG6* | 15 | chr12 | 6417620 | 5.31E-03 | 0.9984 | 1.24E-03 |
| *PODNL1* | 10 | chr19 | 14042840 | 6.26E-03 | 0.9984 | 9.54E-02 |
| *NOVA2* | 16 | chr19 | 46437825 | 6.52E-03 | 0.9984 | 6.94E-02 |
| *HCG27* | 12 | chr6 | 31164776 | 8.07E-03 | 0.9984 | 6.08E-01 |
| *UCKL1* | 15 | chr20 | 62569366 | 8.66E-03 | 0.9984 | 5.09E-01 |
| *TNNI1* | 10 | chr1 | 201371245 | 8.85E-03 | 0.9984 | 2.57E-02 |

*XM stands for the association between exposure to gun violence and methylation.

Table S2. The gene-based methylation mediated effect (IE) on the effect of exposure to gun violence on total IgE change (i.e., gun → methylation → IgE) from top 20 association pairs between exposure to gun violence and gene-based methylation (i.e., gun → methylation).

| Gene | Size | Chr | Position | *P*_XM* | *FDR*_XM | *P*_IE |
|---|---|---|---|---|---|---|
| *ZC3H3* | 63 | chr8 | 144517642 | 7.15E-05 | 0.3128 | 5.04E-02 |
| *DZIP1* | 10 | chr13 | 96234082 | 1.09E-04 | 0.3128 | 6.64E-01 |
| *KIAA0556* | 23 | chr16 | 27564888 | 5.32E-04 | 0.7737 | 7.37E-01 |
| *CDK6* | 22 | chr7 | 92237896 | 6.96E-04 | 0.7737 | 2.30E-01 |
| *LINC00461* | 48 | chr5 | 87835928 | 7.23E-04 | 0.7737 | 8.61E-01 |
| *NOM1* | 18 | chr7 | 156737581 | 8.77E-04 | 0.7737 | 3.58E-01 |
| *ABCA13* | 12 | chr7 | 48237515 | 9.47E-04 | 0.7737 | 7.56E-01 |
| *DACT2* | 17 | chr6 | 168702788 | 1.15E-03 | 0.8256 | 2.40E-01 |
| *TRPV4* | 13 | chr12 | 110220970 | 1.46E-03 | 0.8372 | 1.25E-01 |
| *KBTBD11* | 25 | chr8 | 1919236 | 1.57E-03 | 0.8372 | 7.91E-01 |
| *CRB1* | 10 | chr1 | 197173499 | 1.72E-03 | 0.8372 | 1.41E-01 |
| *SELO* | 10 | chr22 | 50644361 | 1.91E-03 | 0.8372 | 8.99E-01 |
| *FAM124A* | 15 | chr13 | 51792702 | 2.21E-03 | 0.8372 | 9.11E-01 |
| *ACVR1C* | 10 | chr2 | 158451668 | 2.32E-03 | 0.8372 | 9.73E-01 |
| *KANK3* | 18 | chr19 | 8387654 | 2.33E-03 | 0.8372 | 2.03E-01 |
| *LINC01197* | 14 | chr15 | 95821441 | 2.54E-03 | 0.8372 | 8.19E-01 |
| *C2orf48* | 14 | chr2 | 10281159 | 2.62E-03 | 0.8372 | 3.26E-01 |
| *ACTN1* | 22 | chr14 | 69337590 | 2.77E-03 | 0.8372 | 1.58E-01 |
| *PIP5K1C* | 44 | chr19 | 3630649 | 2.78E-03 | 0.8372 | 4.33E-03 |
| *KIAA1804* | 10 | chr1 | 233462711 | 3.08E-03 | 0.8396 | 8.53E-01 |

*XM stands for the association between exposure to gun violence and methylation.
